# The *C. elegans* nervous system reads the internal state of the hydrogen peroxide-detoxification machinery to trigger escape from this common reactive chemical

**DOI:** 10.64898/2026.01.22.701078

**Authors:** Yuyan Xu, Sahana Gangadharan, Maedeh Seyedolmohadesin, Eyob Gebeyaw, Alyson Fulton, Dante Ashih, Meagan Duncan, Matea Zelich, Jun Liu, Mahdi Torkashvand, Daniel Shaw, Katerina Gusarova, Meha Macwan, Avery Kelly, Isidora Beslic, Aishwarya Sood, Olivia Liu, Monika Scholz, Vivek Venkatachalam, Javier Apfeld

**Affiliations:** Biology Department, Northeastern University, Boston, Massachusetts, United States; Bioengineering Department, Northeastern University, Boston, Massachusetts, United States; Physics Department, Northeastern University, Boston, Massachusetts, United States; Max Planck Research Group Neural Information Flow, Max Planck Institute for Neurobiology of Behavior, Bonn, Germany; Chemical Engineering Department, Northeastern University, Boston, Massachusetts, United States

## Abstract

Hydrogen peroxide (H_2_O_2_) is the most common reactive chemical threat faced by organisms. Here, we map the neural circuit that drives chemotactic escape from environmental H_2_O_2_ in the nematode *C. elegans*. Twenty-four neuron classes with sensory endings at the mouth and nose of the animal detect H_2_O_2_. Their response dynamics encode stimulus intensity and exposure history, and their partial redundancy makes avoidance resilient to the loss of individual inputs. Sensing begins when H_2_O_2_ oxidizes the peroxidatic and resolving cysteines of the cytosolic peroxiredoxin PRDX-2, which relays this oxidative signal to cysteines on the LITE-1 and GUR-3 ion channels, triggering calcium influx in sensory neurons that drive escape. Most of these neurons release glutamate to drive H_2_O_2_-dependent excitation of AIA interneurons, whereas others signal through non-glutamatergic routes, providing multiple routes for signal transmission. Thus, the *C. elegans* nervous system acts as a hydrogen peroxide sentinel that monitors H_2_O_2_-induced changes in the intracellular H_2_O_2_-detoxification machinery and relays them to interneurons driving organism-wide escape. This raises the possibility that circuit defects in aging and neurodegenerative disease arise from altered peroxiredoxin-mediated H_2_O_2_ signaling rather than primarily from direct macromolecular damage.

## Introduction

Harmful chemicals threaten organisms by triggering chemical reactions that damage essential cellular macromolecules. Organisms, however, are not passive targets. They have evolved sophisticated defenses that detect, detoxify, and repair such damage. Less understood are the behavioral strategies organisms use to avoid common harmful chemicals in the first place. In this study, we use the nematode *C. elegans* as a model system to investigate how animals sense and escape the most widespread harmful chemical encountered by all organisms: hydrogen peroxide^1–3^.

Hydrogen peroxide (H_2_O_2_) has been the most common reactive chemical threat to life forms since the Great Oxygenation Event 2.5 billion years ago, when cyanobacteria began releasing molecular oxygen into Earth’s atmosphere through photosynthesis^3^. As a result, H_2_O_2_ is produced continuously through the reduction of O_2_ by iron cofactors in numerous enzymes inside cells, and through photochemical reactions both within cells and in the environment^1–3^.

Many organisms—including bacteria, fungi, plants, and animals—also actively produce H_2_O_2_ as a chemical weapon, secreting it at millimolar concentrations to attack prey, competitors, and pathogens^4–6^. While cells employ multiple conserved enzyme systems to degrade H_2_O_2_ and prevent macromolecular damage^1,6^, the mechanisms by which cells—particularly within the nervous system—sense and communicate information about H_2_O_2_ levels remain poorly understood, despite the crucial role of these sensory mechanisms in driving escape behaviors.

Even though hydrogen peroxide is harmful and widespread, organisms use it as a signaling molecule to convey information within cells and across tissues^7^. Compared to other reactive oxygen species (ROS), H_2_O_2_ is more persistent and pervasive due to its nonradical nature, moderate reactivity, and membrane permeability^6^. These properties allow it to travel hundreds of microns from its point of origin^8^, selectively oxidizing sensitive substrates, such as specific protein cysteine thiols, to modify the activity of target proteins^7^. This signaling function is key to various cellular and intercellular processes in both health and disease^7^. Notably, H_2_O_2_ has recently emerged as an important endogenous regulator of nervous system function, influencing neuropeptide and neurotransmitter secretion^9–11^, synaptic plasticity^12,13^, long-term memory formation^14^, and sleep^15^. These observations raise two key questions: What are the molecular mechanisms that enable neurons to detect changes in H_2_O_2_ concentration and convert them into electrical or biochemical signals? And how are these H_2_O_2_-dependent changes in neuronal state propagated and integrated across cells and tissues to coordinate organism-wide responses?

*C. elegans* frequently encounter damaging levels of H_2_O_2_ in their natural environment of rotting fruits and plant stems. H_2_O_2_ produced by *Neorhizobium* sp., a bacterium in the *C. elegans* microbiome, causes DNA damage to the nematode^16^. Pathogenic bacteria such as *Streptococcus pyogenes*, *S. pneumoniae*, *S. oralis*, and *Enterococcus faecium* generate millimolar concentrations of H_2_O_2_ that are lethal to the nematode^17–19^. Environmental sources like plant tissue and rainwater also contain relatively high H_2_O_2_ levels, typically ranging from 10 to 250 µM^20^. In response to H_2_O_2_, *C. elegans* exhibits several aversive behaviors. Recently, we found that escape behavior, which involves coordinated whole-body movement, is mediated at least in part by the ASJ sensory neurons in the amphid chemosensory organs at the nose, which respond strongly to lethal H_2_O_2_ levels^21^. Low levels of H_2_O_2_ and ultraviolet (UV) light also inhibit feeding and promote spitting by activating a neural circuit in the pharyngeal mouth region, via the light-activated ion channels LITE-1 and GUR-3 and the H_2_O_2_-degrading peroxiredoxin PRDX-2^22–26^. The pharyngeal nervous system is largely isolated from the rest of the nervous system, connected only through a single neuron pair^27^. While sensory neurons in the nose are well known to drive chemotaxis^28^, the pharyngeal nervous system had not, to our knowledge, been implicated in regulating whole-body locomotory attraction or repulsion [27]. A better understanding of how different sensory circuits drive escape from H_2_O_2_ in *C. elegans* may provide a framework for studying how more complex animals mount behavioral responses to widespread harmful chemicals.

Here, we map the circuit that drives *C. elegans* chemotactic escape from environmental hydrogen peroxide and uncover the molecular mechanisms by which neurons sense H_2_O_2_ and transmit this information to interneurons controlling locomotory behavior. We identify 24 amphid, labial, and pharyngeal neuron classes projecting to the nose and mouth that detect H_2_O_2_, with response dynamics that encode stimulus intensity and exposure history; the partial redundancy of many of these neurons in promoting H_2_O_2_ avoidance confers circuit-level resilience to input failures. Most of these neurons release glutamate, which is required for H_2_O_2_-dependent excitation of the locomotion-regulating AIA interneurons, yet some use non-glutamatergic routes, ensuring transmission fault tolerance. H_2_O_2_ sensing begins when it oxidizes the peroxidatic (C55) and resolving (C176) cysteines of the cytosolic peroxiredoxin PRDX-2, enabling the enzyme to serve a dual role: detoxifying peroxide and conveying its presence. The thioredoxin system, comprising the NADPH → TRXR-1 → TRX-1 (C38/C72) redox relay, acts as a molecular wire that returns electrons to oxidized PRDX-2, resetting the sensor. In each cycle, oxidized PRDX-2 can transfer its oxidative equivalent to specific cytosol-facing cysteines on the LITE-1 and GUR-3 ion channels, triggering calcium influx in specific nose and mouth sensory neurons that drive chemotactic escape. Thus, the *C. elegans* nervous system functions as a hydrogen peroxide sentinel, with sensory neurons converting the internal state of the detoxifying machinery of this harmful chemical into actions that protect the whole organism.

## Results

### *C. elegans* chemotaxes away from non-lethal concentrations of H_2_O_2_

In their natural environment, *C. elegans* nematodes may encounter harmful hydrogen peroxide levels due to the presence of H_2_O_2_-producing bacteria in their microbiome and surrounding habitat^16,21^. These bacteria can release millimolar concentrations of H_2_O_2_ ^17–19^. Previously, we found that *C. elegans* hermaphrodites exhibit an aversive locomotory response—reversing direction followed by performing an omega bend^29–31^—when they encounter a drop of 1 mM H_2_O_2_ ^21^, a concentration that is lethal to embryos, L1 larvae, and adults^18,19,21,32,33^. Here, we set out to determine the extent to which *C. elegans* can navigate away from a concentration gradient of hydrogen peroxide.

We observed the migration of populations of day 1 adult nematodes placed at the center of a 9 cm petri plate, exposed to a radial Gaussian H_2_O_2_ gradient originating from a point source 3 cm away (Figure 1A-B). The H_2_O_2_ concentration was estimated to be 6 mM at the source, 0.12 mM at the plate center, and 0.4 µM at a distance of 6 cm from the source (Figure 1A). Within 10 min, animals vacated regions estimated to exceed 0.1 mM H_2_O_2_, while those that remained in these higher-concentration zones appeared to move more rapidly (Figure 1B, Supplementary Video 2). These avoidance behaviors were not observed in the absence of H_2_O_2_ (Figure 1B, Supplementary Video 1).

**Figure 1.**
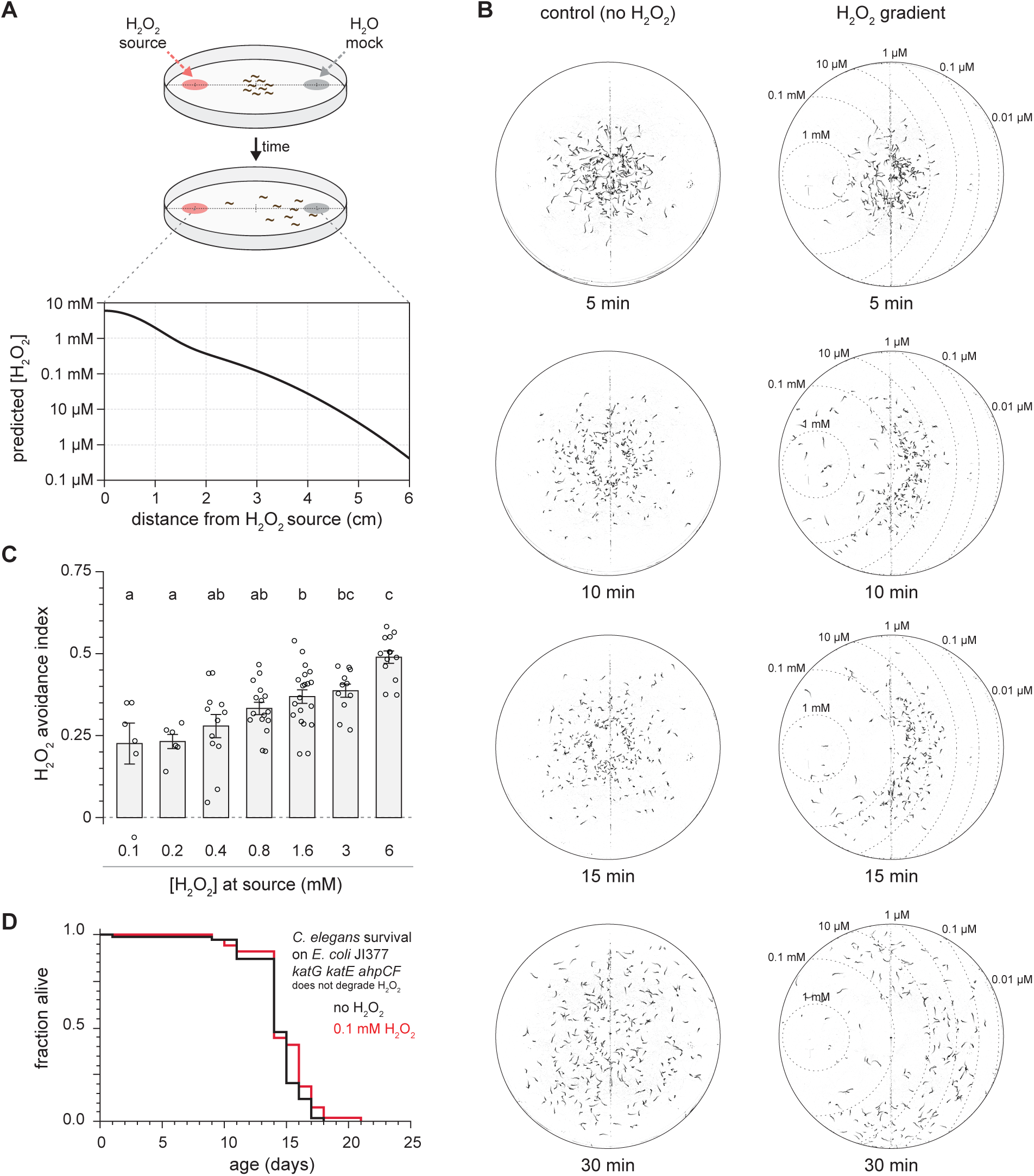
*C. elegans* chemotaxes away from non-lethal concentrations of H_2_O_2_. (A) Schematic of the chemotaxis assay (top) and the estimated H_2_O_2_ concentration along the gradient originating from the point source (bottom). (B) Time-series images showing 20-second trajectories of individual worms at the indicated time points on plates without added H_2_O_2_ (left) and with 6 mM H_2_O_2_ at the point source (right). Dotted lines mark estimated H_2_O_2_ isoconcentration contours. (C) H_2_O_2_ avoidance changed with concentration. Data are plotted as mean ± s.e.m. Groups labeled with different letters exhibited significant differences (*P* < 0.05, Tukey HSD test), otherwise not significant (*P* > 0.05). (D) *C. elegans* lifespan was not affected by 0.1 mM H_2_O_2_ (*P* > 0.05 log-rank test). Worms grown on *E. coli* JI377, which does not degrade environmental H_2_O_2_.

To quantify how sensitively *C. elegans* avoids H_2_O_2_, we measured chemotactic escape as a function of H_2_O_2_ concentration using a binary choice assay^34^. Before the assay, we applied the paralytic agent sodium azide at both the H_2_O_2_ source and a control spot on the opposite side of the petri plate to immobilize nematodes that reached those locations (Figure S1A). After 2 hours, we recorded the nematodes’ final positions (Figure S1B). An avoidance index of –1 indicated complete preference for the H_2_O_2_ source area, +1 indicated complete preference for the control spot area, and 0 indicated no preference^35^. *C. elegans* avoided H_2_O_2_ across a wide range of concentrations (Figure 1C). L4 larvae already showed a robust avoidance of 6 mM H_2_O_2_, while adult worms, from the onset of adulthood to day 2, maintained a steady, slightly increased level of avoidance (Figure S1C). In addition, avoidance of 6 mM H_2_O_2_ by day 1 adult worms isolated and assayed in complete darkness was indistinguishable from that observed under normal lab lighting conditions (Figure S1D), indicating that their behavioral response to H_2_O_2_ does not depend on light. H_2_O_2_ avoidance was dose-dependent, with an index of 0.23 with 0.1 mM H_2_O_2_ at the source and progressively increasing up to 0.49 with 6 mM H_2_O_2_ at the source (Figure 1C).

Since *C. elegans* adults avoided H_2_O_2_ gradients with a maximum concentration of 0.1 mM, we investigated whether this concentration affected their survival during adulthood. Exposure to 0.1 mM H_2_O_2_ did not affect the lifespan of nematodes grown on *E. coli* JI377. This strain, a *katG katE ahpCF* triple null mutant, is unable to degrade environmental H_2_O_2_, thus allowing us to raise the worms in a constant H_2_O_2_ environment^21,32,33,36^ (Figure 1D). These findings indicated that *C. elegans* can chemotax away even from concentrations of hydrogen peroxide that are not visibly deleterious to their survival.

### Sensory neurons in both the somatic and pharyngeal nervous systems promote chemotactic avoidance of H_2_O_2_

We set out to identify which neurons are required for chemotactic avoidance of hydrogen peroxide. Initially, we focused on ciliated sensory neurons, which enable nematodes to detect chemical and physical stimuli (Figure 2A)^34,37–40^. Using a binary choice assay, we measured avoidance from a 6 mM H_2_O_2_ source in mutants with severe sensory defects. Two cilium-structure mutants, *osm-5* and *osm-6*, which have severely truncated sensory endings in most ciliated sensory neurons due to impairment of intraflagellar transport^41^, showed a 0.2 reduction in avoidance index compared to wild-type controls (Figure 2B), indicating a role for ciliated sensory neurons in promoting H_2_O_2_ avoidance.

**Figure 2.**
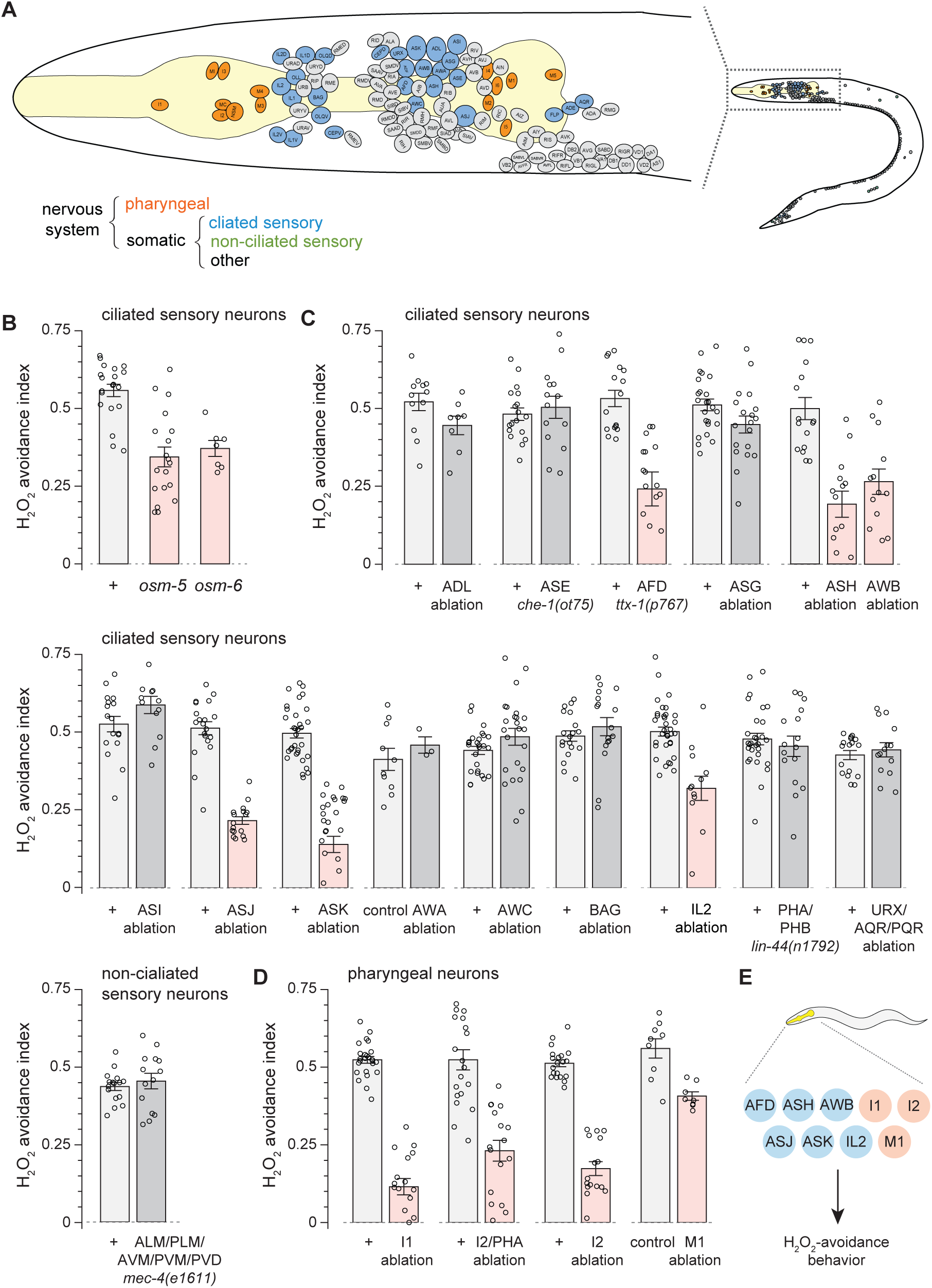
Sensory neurons in both the somatic and pharyngeal nervous systems promote chemotactic avoidance of H_2_O_2_. (A) Diagram of the *C. elegans* hermaphrodite nervous system showing the positions of neuronal nuclei (modified from Ref. ^95^ with permission). The pharynx (yellow) is the worm’s feeding organ. Selected subsets of neurons are colored. (B-D) H_2_O_2_ avoidance of nematodes with defects in sensory cilia (B), or with genetic ablation of specific ciliated (C) or pharyngeal (D) neurons. Data are plotted as mean ± s.e.m. Groups labeled with a red bar exhibited significant differences relative to their respective wild-type control (*P* < 0.05, ANOVA), otherwise not significant (*P* > 0.05). (E) Specific somatic (blue) and pharyngeal (orange) neuronal classes promote H_2_O_2_ chemotactic avoidance.

That said, we were surprised that cilium-structure mutants retained a substantial ability to avoid H_2_O_2_, despite being presumed to lack all sensory responses mediated by ciliated sensory neurons^41–44^. Recent studies showed that in *osm-6* (and other cilium-structure mutants) some neurons retain the capacity to respond to certain odorants^45,46^. This residual activity could explain the remaining behavioral avoidance of H_2_O_2_ in *osm-5* and *osm-6* mutants. Alternatively, cilium-independent processes within ciliated neurons, non-ciliated neurons, or other cell types might also contribute to this residual avoidance behavior. The *C. elegans* hermaphrodite nervous system has two largely synaptically isolated components: the somatic nervous system (282 neurons, including 62 ciliated and 10 non-ciliated sensory neurons) and the pharyngeal nervous system (20 non-ciliated neurons)^27,40,47,48^ (Figure 2A). Although previously thought to lack sensory neurons, recent ultrastructural analysis revealed that most pharyngeal neurons likely have sensory functions^27^. While the pharyngeal nervous system has not been previously implicated in chemotaxis, the I2 pharyngeal neuron pair was found to sense H_2_O_2_^22^, raising the possibility that both pharyngeal and somatic sensory neurons might jointly regulate H_2_O_2_ chemotactic avoidance.

To identify neurons involved in hydrogen peroxide avoidance, we systematically quantified H_2_O_2_ avoidance in a collection of strains in which specific classes of sensory neurons were genetically ablated using neuron-specific expression of caspases^32,49–57^, mutations affecting neuron-specific fate determinants^58,59^, or a mutation causing degeneration of specific touch-receptor neurons^60^. We also examined a strain in which defective tail socket cell positioning prevented specific tail sensory neurons from being directly exposed to the external environment^61,62^. Because we could not identify a single promoter expressed exclusively in I2 neurons, we initially ablated I2 using *flp-15* driven caspase, which also targeted PHA. To achieve I2 specificity, we then used the split cGAL intersectional system^63^, using the *aqp-5* promoter to express cGAL-N in I2 and M3 and the *gur-3* promoter to express cGAL-C in I2 and I4; this approach reconstituted a functional cGAL driver exclusively in I2, driving their selective ablation in animals carrying a *UAS::caspase* effector transgene. Our collection covered 22 of the 32 classes of sensory neurons in the somatic nervous system, including 11 out of the 12 classes of ciliated neurons that make up the two amphids (the major sensory organs), 7 of the 14 classes of non-amphid ciliated neurons, and 4 of the 6 classes of non-ciliated sensory neurons (Figure 2A, Table S1). Within the pharyngeal nervous system, we covered 3 of the 11 classes of known or likely sensory neurons^24,27^.

Individual ablation of four amphid sensory neuron classes (ASH, ASJ, ASK, AWB), one non-amphid ciliated neuron class (IL2), and three pharyngeal sensory neuron classes (I1, I2, M1) significantly reduced, but did not eliminate, avoidance from a 6 mM H_2_O_2_ source (Figures 2C-D). Additionally, H_2_O_2_ avoidance was reduced by a mutation in *ttx-1* (Figure 2C), which blocks microvilli formation within the AFD sensory cilium^41,59^. In contrast, ablation of the remaining neuron classes, including ADL, ASE, ASG, ASI, AWA, AWC, BAG, joint ablation of URX, AQR, and PQR, and of ALM, PLM, AVM, PVM, and PVD, and misplacement of the socket cell connecting PHA and PHB to the environment, had no effect on H_2_O_2_ avoidance (Figure 2C). Taken together, we identified six somatic and three pharyngeal sensory neuron classes that promote H_2_O_2_ chemotactic avoidance (Figure 2E). Because these neuronal classes contribute to but are not individually essential for complete H_2_O_2_ avoidance behavior, we infer that they function in a partially redundant manner. Their redundancy suggests a distributed sensory neuronal strategy that ensures the robustness of the behavior to the loss of individual sensory classes.

### Sensory neurons in both the somatic and pharyngeal nervous systems exhibit neuron-specific, dose-dependent responses to H_2_O_2_

Sensory neurons in the somatic and pharyngeal nervous systems could promote hydrogen peroxide avoidance by directly detecting H_2_O_2_ or by acting in parallel or downstream of its perception. Their responses may differ in sensitivity, dynamic range, and dependence on prior exposure. To clarify these roles, we monitored their activity in response to a wide range of H_2_O_2_ concentrations.

The activity of *C. elegans* sensory neurons strongly correlates with their calcium dynamics^64^. To investigate their responses to H_2_O_2_, we imaged fluorescence of the calcium indicator GCaMP6 expressed in the nuclei of either all neurons or only ciliated sensory neurons. We imaged responses to H_2_O_2_ pulses across a broad range of concentrations: typically 10 µM to 10 mM, but extended in some trials from as low as 100 nM to as high as 100 mM. We delivered H_2_O_2_ using a custom-built microfluidic device that allows odorants to flow past the nose and mouth of an immobilized day 1 adult (Figure 3A-D)^21,65^. We provided increasing concentrations of H_2_O_2_, with 15-second exposure intervals preceded and followed by 45-second buffer (H_2_O) intervals. We recorded the activity from the nuclei of 30 somatic sensory neurons projecting to the amphid and labial organs in the nose, and all 20 neurons in the pharyngeal nervous system (Figure 3D). Our imaging studies covered all 12 amphid neuron classes (ADF, ADL, AFD, ASE, ASG, ASH, ASI, ASJ, ASK, AWA, AWB, and AWC), 3 labial neuron classes (BAG, IL2V, and URX), and all 14 pharyngeal neuron classes (I1, I2, I3, I4, I5, I6, M1, M2, M3, M4, M5, MC, MI, and NSM).

**Figure 3.**
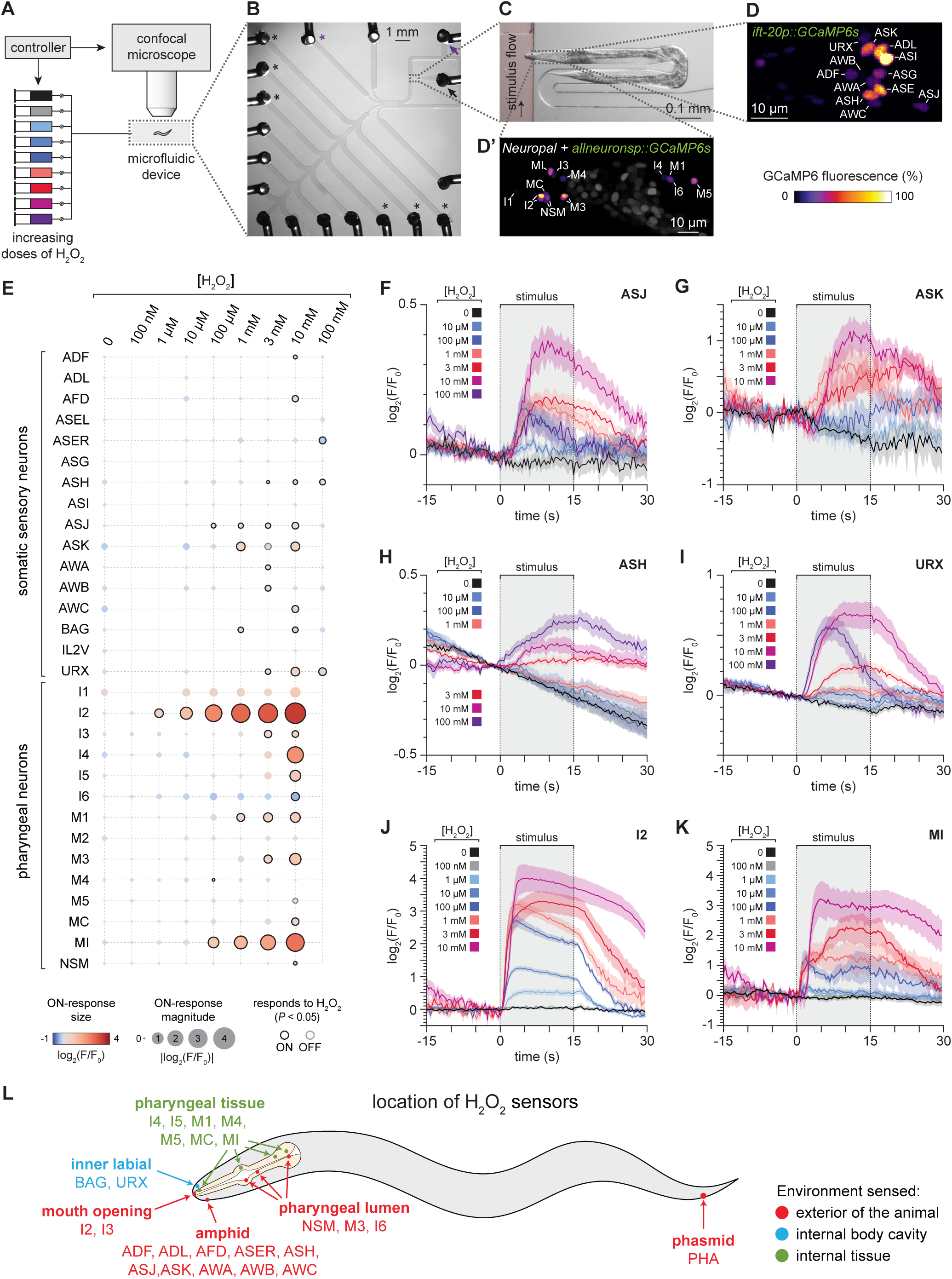
Sensory neurons in both the somatic and pharyngeal nervous systems exhibit neuron-specific, dose-dependent responses to H_2_O_2_. (A) Schematic of the microfluidic platform used for controlled stimulus delivery and calcium imaging. (B) Image of the multi-channel device. Black arrow: worm-loading inlet; purple arrow: waste outlet. Black asterisks: stimulus delivery channels used; purple asterisk: buffer channel. (C) Enlarged view of the channel where the sensory endings in the nose and mouth of the immobilized nematode are stimulated with H_2_O_2_. (D-D’) Relative GCaMP6 fluorescence from head neurons imaged in worms expressing GCaMP6s (D) in ciliated neurons (12 classes shown) or (D’) all neurons (12 pharyngeal classes colored), imaged at single-cell resolution. (E) Responses of somatic and pharyngeal neuron classes to specific concentrations of H_2_O_2_. Each circle represents one neuron class at a given H_2_O_2_ concentration. Circle color encodes the mean change in GCaMP6 fluorescence during the 15-second stimulus interval (ON response); circle area scales with the absolute response magnitude. Black rings indicate ON responses that differ significantly from the no-H_2_O_2_ control (Dunnett’s test, *P* < 0.05); gray rings indicate significant OFF-responses measured during the 15-second post-stimulus interval (Dunnett’s test, *P* < 0.05); unmarked circles indicate responses that were not significant in either interval (*P* > 0.05). (F-K) Mean GCaMP6 fluorescence traces for (F) ASJ, (G) ASK, (H) ASH, (I) URX, (J) I2, and (K) MI neuronal classes in response to the indicated H_2_O_2_ concentrations. Data are plotted as mean ± s.e.m. The shaded box marks the stimulus delivery interval. The remaining neuronal classes are shown in Figure S2. (L) Schematic illustrating the location of the sensory endings of H_2_O_2_-sensing neurons. Endings sample distinct environments, positioning the network to detect H_2_O_2_ from external and internal sources. Statistical analyses for panels (E-K) are in Table S6.

Hydrogen peroxide modulated the activity of 24 of 29 neuronal classes examined: it increased [Ca^2+^] in 21 classes—ten somatic (ADF, ADL, AFD, ASH, ASJ, ASK, AWA, AWB, AWC, URX) and eleven pharyngeal (I2, I3, I4, I5, M1, M3, M4, M5, MC, MI, NSM)—and decreased [Ca^2+^] in 3 classes—two somatic (ASER, BAG) and one pharyngeal (I6) (Figure 3E-K, Figure S2). Under identical illumination conditions, recordings without exogenous H_2_O_2_ showed no calcium responses, consistent with and extending our previous studies^21,66^, confirming that the observed responses represent H_2_O_2_-evoked activity that is not evoked by illumination alone. These findings are consistent, and substantially extend, previous studies by our labs and others showing that the ASJ, ASK, URX, and I2 neuron classes respond to 1 mM H_2_O_2_^21,25^. The 24 H_2_O_2_-responsive neuronal classes map to the amphid and labial sensory organs in the nose, as well as the mouth opening, corpus, sieve, isthmus, grinder, and terminal bulb in the pharynx (Figure 3L). In the nose, every H_2_O_2_-sensing neuron we identified contacts the exterior of the animal, except for BAG and URX, whose endings face the pseudocoelomic body cavity^67^. In the pharynx, I2, I3, I6, M3, and NSM have sensory endings exposed to the exterior lumen, while the remaining classes are exposed to distinct internal compartments^27^. Thus, H_2_O_2_-sensing neurons are distributed across multiple anatomical compartments and are positioned to detect both external and internal sources of hydrogen peroxide.

H_2_O_2_-evoked activity varied with both neuron class and stimulus strength. I2 neurons at the mouth opening displayed a graded response across 1 µM-10 mM, with amplitudes rising with concentration (Figure 3J). ASH, ASJ, URX, and MI showed similar gradation, whereas all other classes appeared to respond in a discrete, all-or-none manner (Figures 3F-K; Figure S2).

H_2_O_2_-detection thresholds differed widely across neuron classes (Figure 3E). I2 was the most sensitive, responding at 1 µM (Figure 3J). ASJ, M4, and MI responded at 100 µM; ASK, BAG, and M1 at 1 mM; ASH, AWA, AWB, URX, I3, and M3 at 3 mM; ADF, AFD, AWC, I4, I5, I6, M5, MC, and NSM at 10 mM; and ASER only at 100 mM. AFD, ADL, AWB, AWC, BAG, and M4 responded inconsistently, exhibiting H_2_O_2_-evoked activity in only a subset of experiments, making it difficult to determine their detection threshold. Because sensory endings reside in distinct compartments (Figure 3L), the H_2_O_2_ concentration they experienced may differ from the delivered dose. In addition, because AFD activity is highly sensitive to temperature^64,68^, we cannot rule out that their response to 10 mM H_2_O_2_ were due to slight temperature differences between solutions rather than H_2_O_2_ itself.

H_2_O_2_-response dynamics also varied across neuron classes at both stimulus onset (ON responses) and stimulus removal (OFF responses) (Figures 3F-K; Figure S2). Some classes responded rapidly upon H_2_O_2_ exposure (I2, MI, ASJ, AWC, URX), while others showed slower ON responses (ADF, ASER, ASK, ASH, I3, I4, I5, M1, M3, M5). Similarly, rapid OFF responses occurred in certain classes (I2, I3, I4, AWC, URX), whereas others desensitized more slowly (ASER, ASH, ASJ, ASK, AWA, AWB, URX, I5, M1, M3, M5, MI).

Finally, response dynamics often depended on both H_2_O_2_ concentration and exposure history. In I2 neurons, 1 µM H_2_O_2_ evoked a rapid, new steady-state response followed by an exponentially decaying OFF response (Figure 3J). At 10 µM, the ON response overshot before settling at a lower steady state. At still higher concentrations, a steady state was not reached, and the onset of the recovery phase was delayed relative to stimulus removal. Occasionally, a 1 mM H_2_O_2_ stimulus either abolished recovery during the OFF response—rendering the neuron unresponsive to later 3 mM and 10 mM stimuli—or only partially terminated it, markedly blunting subsequent responses (Figure S3A). Similar desensitization occasionally occurred in ASJ (Figure S3B), M1, M3, and MI, underscoring that many neuronal classes integrate both hydrogen peroxide concentration and exposure history.

The 24 neuronal classes exhibiting H_2_O_2_-induced changes in activity included all but two (I1, IL2) of the neuronal classes that promoted H_2_O_2_ chemotactic avoidance: AFD, ASH, ASJ, ASK, and AWB in the somatic nervous system, and I2 and M1 in the pharyngeal nervous system. However, several neuronal classes that responded to H_2_O_2_, including ADL, ASE, AWC, BAG, and URX, or that were known to respond, namely PHA^25^, did not affect H_2_O_2_ chemotactic avoidance. These neurons may be necessary for H_2_O_2_ avoidance in other contexts or mediate other H_2_O_2_-dependent physiological responses. Taken together, these findings suggested that multiple neuronal classes in both the somatic and pharyngeal nervous systems promoted hydrogen peroxide avoidance upon sensing this chemical.

### The cytosolic peroxiredoxins PRDX-2 and PRDX-6 promote chemotactic avoidance of H_2_O_2_

To understand the mechanisms by which hydrogen peroxide triggered avoidance, we first examined the behavior of mutants with impaired intracellular H_2_O_2_ degradation. We focused on peroxiredoxins (PRXs), which are the major enzymes responsible for rapidly degrading H_2_O_2_ at low concentrations (<10 µM)^1^. The *C. elegans* genome contains two 2-Cys PRX-coding genes, *prdx-2* and *prdx-3*, which encode cytosolic and mitochondrial PRXs, respectively^69,70^, as well as a single 1-Cys PRX-coding gene, *prdx-6*, which encodes a cytosolic protein. We hypothesized that these mutants would show an increase in H_2_O_2_ avoidance because, due to defective degradation, *prdx-2* and *prdx-3* mutants have higher intracellular H_2_O_2_ concentrations^9,71^. Contrary to our expectation, both *prdx-2(gk169)* null mutants and *prdx-6(tm4225)* null mutants showed a 0.3 reduction in H_2_O_2_ avoidance index compared to wild-type controls (Figure 4A), while *prdx-3(gk529)* null mutants did not affect H_2_O_2_ avoidance (Figure 4A). These findings suggested that, rather than merely degrading H_2_O_2_, the cytosolic peroxiredoxins PRDX-2 and PRDX-6 actively facilitate H_2_O_2_ avoidance.

**Figure 4.**
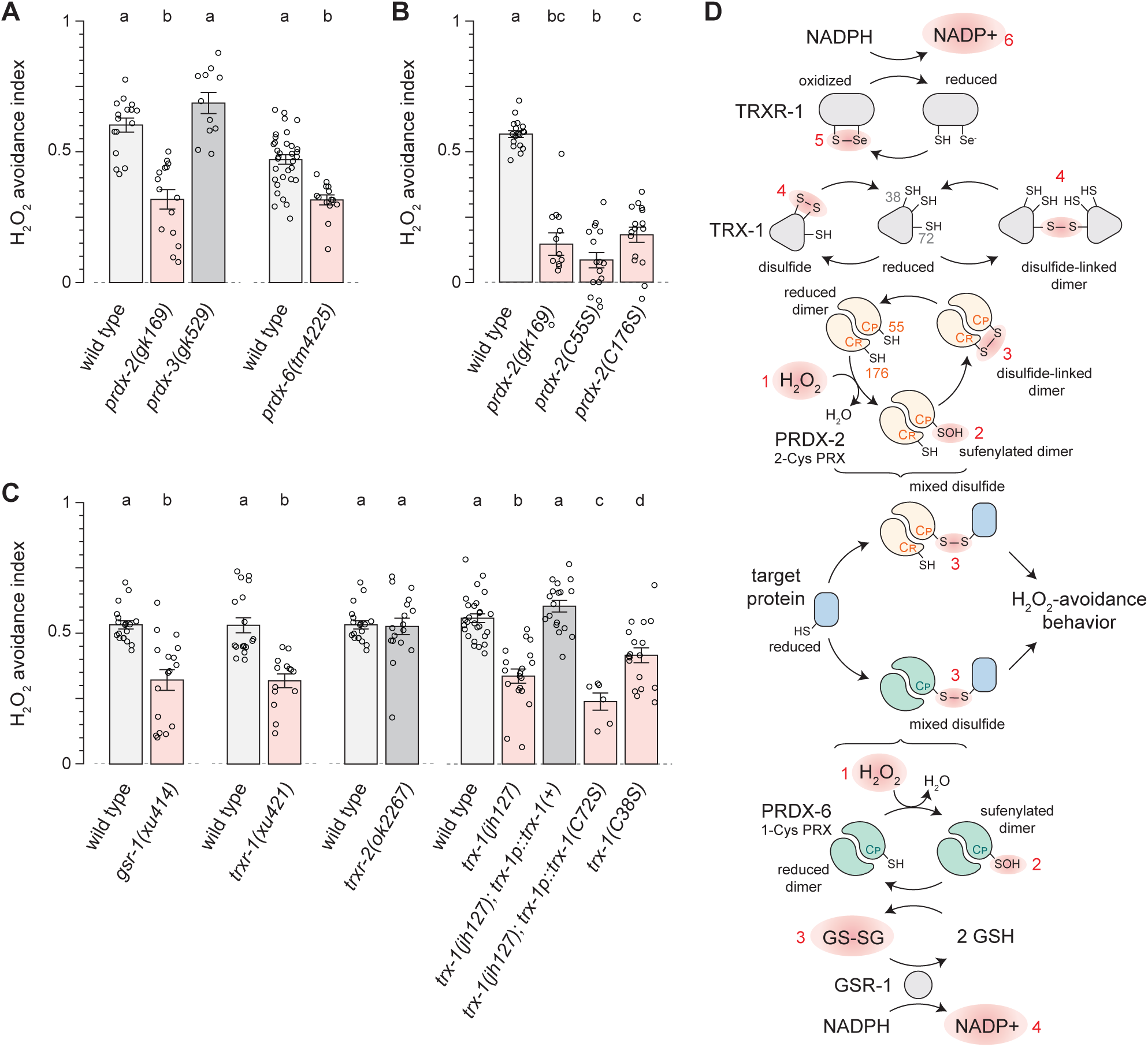
The cytosolic peroxiredoxins PRDX-2 and PRDX-6 promote chemotactic avoidance of H_2_O_2_. (A) H_2_O_2_ avoidance was lowered in mutants lacking cytosolic *prdx-2* and *prdx-6,* but was unaffected by loss of mitochondrial *prdx-3*. (B) Replacing the peroxidatic (C55) or resolving (C176) cysteine of *prdx-2* with serine lowered H_2_O_2_ avoidance to *prdx-2* null levels. (C) H_2_O_2_ avoidance was lowered by mutations in the glutathione reductase *gsr-1* and cytosolic thioredoxin reductase *trxr-1*, but not by loss of the mitochondrial *trxr-2*. Loss of the ASJ-specific thioredoxin *trx-1*, or replacement of its catalytic (C38) or non-catalytic (C72) cysteines to serine also lowered H_2_O_2_ avoidance. Data are plotted as mean ± s.e.m. Groups labeled with different letters exhibited significant differences (*P* < 0.05, Tukey HSD test), otherwise not significant (*P* > 0.05). (D) Proposed mechanism linking H_2_O_2_ degradation and chemotactic avoidance. H_2_O_2_ sensing begins when it oxidizes the peroxidatic (C_P_) cysteine of 2-Cys PRDX-2 or 1-Cys PRDX-6. Upon oxidation, these enzymes can relay the presence of H_2_O_2_ by oxidizing target proteins that trigger H_2_O_2_ avoidance behavior. In PRDX-2 the resolving cysteine (C_R_) reacts with oxidized C_P_, forming a disulfide. The thioredoxin system, comprising the NADPH → TRXR-1 → TRX-1 (C38/C72) relay, then acts as a molecular wire that returns electrons to oxidized PRDX-2, resetting the sensor. Other TRX-like proteins may play analogous roles in other neurons to the ASJ-specific function TRX-1. In parallel, the glutathione system, comprising the NADPH → GSR-1 → GSH relay, acts similarly to return electrons to oxidized PRDX-6. We speculate that the resetting of the PRDX-2 and PRDX-6 sensors is needed for H_2_O_2_ avoidance because it prevents them from entering an inactive state. For example, without rapid reduction, additional H_2_O_2_ molecules might sequentially hyperoxidize the peroxidatic cysteines, first to sulfinic acid and then to sulfonic acid, irreversibly disabling the sensor (see Discussion). Red shading and numbered steps trace the sequential transfer of oxidative equivalents between species along the redox relays.

Motivated by these findings, we tested whether H_2_O_2_ avoidance was directly coupled to H_2_O_2_ degradation by PRDX-2. In the first step of the PRX catalytic cycle (Figure 4D), H_2_O_2_ oxidizes the thiol group (–SH) of the peroxidatic cysteine (C_P_) in the enzyme’s active site to a sulfenic acid (–SOH) intermediate, which is then recycled to the reduced state in subsequent steps of the cycle^69^. Using CRISPR, we replaced the codon for this peroxidatic cysteine in the endogenous *prdx-2* gene with serine, an amino acid isosteric with cysteine but with an effectively redox-inactive hydroxyl group instead of a redox-active thiol. The resulting *prdx-2(C55S)* mutation lowered H_2_O_2_ avoidance as much as the null allele (Figure 4B). Therefore, the peroxidatic cysteine is essential for PRDX-2-mediated H_2_O_2_ avoidance. These findings suggested that H_2_O_2_ avoidance is directly coupled to the first step of the catalytic cycle of H_2_O_2_ degradation by PRDX-2.

We next asked whether mechanisms involved in the recycling of the sulfenylated C_P_ back to the reduced state were required for H_2_O_2_ avoidance. These recycling mechanisms differ between 1-Cys and 2-Cys PRXs (Figure 4D). 2-Cys PRXs use a dedicated resolving cysteine to form an inter-subunit disulfide that is recycled by the thioredoxin system^72^, whereas 1-Cys PRXs lack this resolving cysteine and instead rely on glutathione to reduce their sulfenylated peroxidatic cysteine^73,74^. In principle, blocking C_P_ recycling could increase H_2_O_2_ avoidance by prolonging the lifetime of the sulfenylated, signaling-competent PRX species. Alternatively, it could lower H_2_O_2_ avoidance by trapping PRX in a signaling-inactive state or by promoting the formation of additional inactive species through side reactions such as C_P_ hyperoxidation.

Recycling of 1-Cys PRXs depends on the glutathione redox relay: the oxidized C_P_ is sequentially reduced by two glutathione (GSH) molecules, producing glutathione disulfide (GSSG)^73,74^, which is then recycled back to GSH by glutathione reductase (GR) using electrons from NADPH (Figure 4D)^75^. We found that the *xu414* mutant allele in *gsr-1*, which introduces a G279E missense mutation in a conserved site in the FAD binding domain of the sole *C. elegans* glutathione reductase^76–78^, lowered the H_2_O_2_ avoidance index by 0.21 compared to wild-type controls (Figure 4C). We propose that by increasing the GSH/GSSG ratio, GSR-1 facilitates the recycling of PRDX-6 to its active, reduced state, leading to improved H_2_O_2_ avoidance (Figure 4D).

Recycling of 2-Cys PRXs is mediated by a multi-step process involving the thioredoxin redox relay. PRXs function as obligate homodimers^72^. In the first step of recycling of 2-Cys PRXs, the resolving cysteine (C_R_) from the opposing subunit of the PRX dimer reacts with the oxidized peroxidatic cysteine, forming a disulfide bond^72^ (Figure 4D). Consistent with this mechanism, biochemical studies show disulfide-linked PRDX-2 accumulates after *C. elegans* are exposed to 0.2 mM H_2_O_2_ for 5 minutes^69^. Using CRISPR, we replaced the codon for this resolving cysteine in the endogenous *prdx-2* gene with one coding for serine. The resulting *prdx-2(C176S)* mutation lowered H_2_O_2_ avoidance as much as the C55S and null alleles (Figure 4B), indicating that both C_P_ and C_R_ promote H_2_O_2_ avoidance.

Disulfide-bonded PRX dimers are then recycled back to their reduced form by the thioredoxin (TRX)-thioredoxin reductase (TRXR) system^79^: TRX reduces the disulfide-bonded PRX dimer while becoming oxidized itself, and TRXR subsequently restores reduced TRX using electrons from NADPH (Figure 4D)^75^. We found that the *xu421* stop-codon mutation in *trxr-1*^76^, which encodes the sole cytosolic TRXR in the *C. elegans* genome^80,81^, lowered the H_2_O_2_ avoidance index by 0.2 relative to wild type (Figure 4C), while a null mutation in *trxr-2,* which encodes the sole mitochondrial TRXR^81^, had no effect (Figure 4C). Of the at least eight thioredoxins in the *C. elegans* genome, we focused on *trx-1* because it is expressed exclusively in the cytosol of ASJ^82^, an H_2_O_2_-sensing neuron that promoted H_2_O_2_ avoidance (Figures 2C and 3F). *trx-1(jh127)* null mutants had impaired H_2_O_2_ avoidance, and this defect was rescued by a single copy transgene restoring TRX-1 expression only in in ASJ (Figure 4C)^83^. We observed a similar reduction with the *trx-1(nu517)* mutation, which replaces the TRX-1 catalytic cysteine 38 with serine (Figure 4C). Cysteine 72 of TRX-1 is necessary for sensory perception of nitric oxide by ASJ^83^ and is required for TRX dimerization through disulfide bridges^84^. This dimerization masks the catalytic site of TRX, preventing recycling of its catalytic cysteine by TRXR (Figure 4D)^84,85^. A single-copy transgene expressing TRX-1(C72S) in ASJ failed to rescue the H_2_O_2_ avoidance defect of *trx-1(nu517)* (Figure 4D). We conclude that the TRX-1-TRXR-1 system is necessary for H_2_O_2_ avoidance via a mechanism that relies on TRX-1 catalytic C38 and TRXR-1, as well as on TRX-1 C72, which is not involved in catalysis. We propose that the TRX-1-TRXR-1 redox relay promotes H_2_O_2_ avoidance by promoting the recycling of disulfide-bonded PRDX-2 dimers to their reduced form, acting like a wire through which electrons flow sequentially from NADPH to TRXR-1, then to TRX-1 cysteines C38 and C72, and finally to the C_R_ and C_P_ cysteines of PRDX-2 (Figure 4D).

Based on these findings, we propose that sulfenylated peroxiredoxin, generated when the peroxidatic cysteine is oxidized by H_2_O_2_, serves as the central signaling intermediate that couples cytosolic H_2_O_2_ to chemotactic escape (Figure 4D). We favor a model where recycling pathways promote H_2_O_2_ avoidance by regenerating reduced, sensing-competent PRXs and preventing the formation of additional inactive species through further H_2_O_2_-dependent hyperoxidation of C_P._ In this way, peroxiredoxin-based redox wires remain functionally engaged, even though recycling depletes the sulfenylated intermediates that carry the H_2_O_2_ signal. We elaborate on this model in the Discussion.

### Perception of H_2_O_2_ by sensory neurons in both the somatic and pharyngeal nervous systems requires the cytosolic peroxiredoxin PRDX-2

Since *prdx-2* was required for H_2_O_2_ avoidance, we next determined whether it was also necessary for H_2_O_2_-evoked changes in the activity of somatic and pharyngeal sensory neurons. PRDX-2 is highly abundant^86^ and broadly expressed^22,25,70^, and single-neuron mRNA-seq from the CeNGEN consortium showed that all H_2_O_2_-sensing neurons we identified express *prdx-2*, with I2 exhibiting the highest expression (about twenty fold higher than the median neuron)^87^. Using a modified version of our previous calcium imaging setup, we simultaneously imaged wild-type worms and *prdx-2(gk169)* mutants—along with additional mutants described later in the manuscript—while exposing them to either 0 or 10 mM H_2_O_2_. This updated microfluidic design enabled us to image three immobilized worms at the same time and deliver up to five different stimuli (Figure S4), with worms of different genotypes imaged simultaneously. We focused on neurons that showed consistent H_2_O_2_-evoked changes in activity. In *prdx-2(−)* mutants, all H_2_O_2_-responsive pharyngeal neurons exhibited defective or absent responses and most, though not all, H_2_O_2_-responsive somatic sensory neurons were also affected (Figure 5A). In the pharynx, *prdx-2* loss abolished responses in I4, I6, M3, M4, M5, MC, and NSM, severely reduced I2 and M1 responses, and moderately reduced MI responses (Figure 5B, Figure S5). In the nose, it abolished ASK and ASJ responses, while ADF, ADL, ASH, and URX remained unaffected. These findings extend prior work showing that *prdx-2* is partially required for I2 responses to 1 mM and 8.9 M H_2_O_2_ ^22,25^. Thus, PRDX-2 was critical for H_2_O_2_ perception by pharyngeal (I2, M1) and somatic (ASK, ASJ) neuronal classes that promoted avoidance of this chemical.

**Figure 5.**
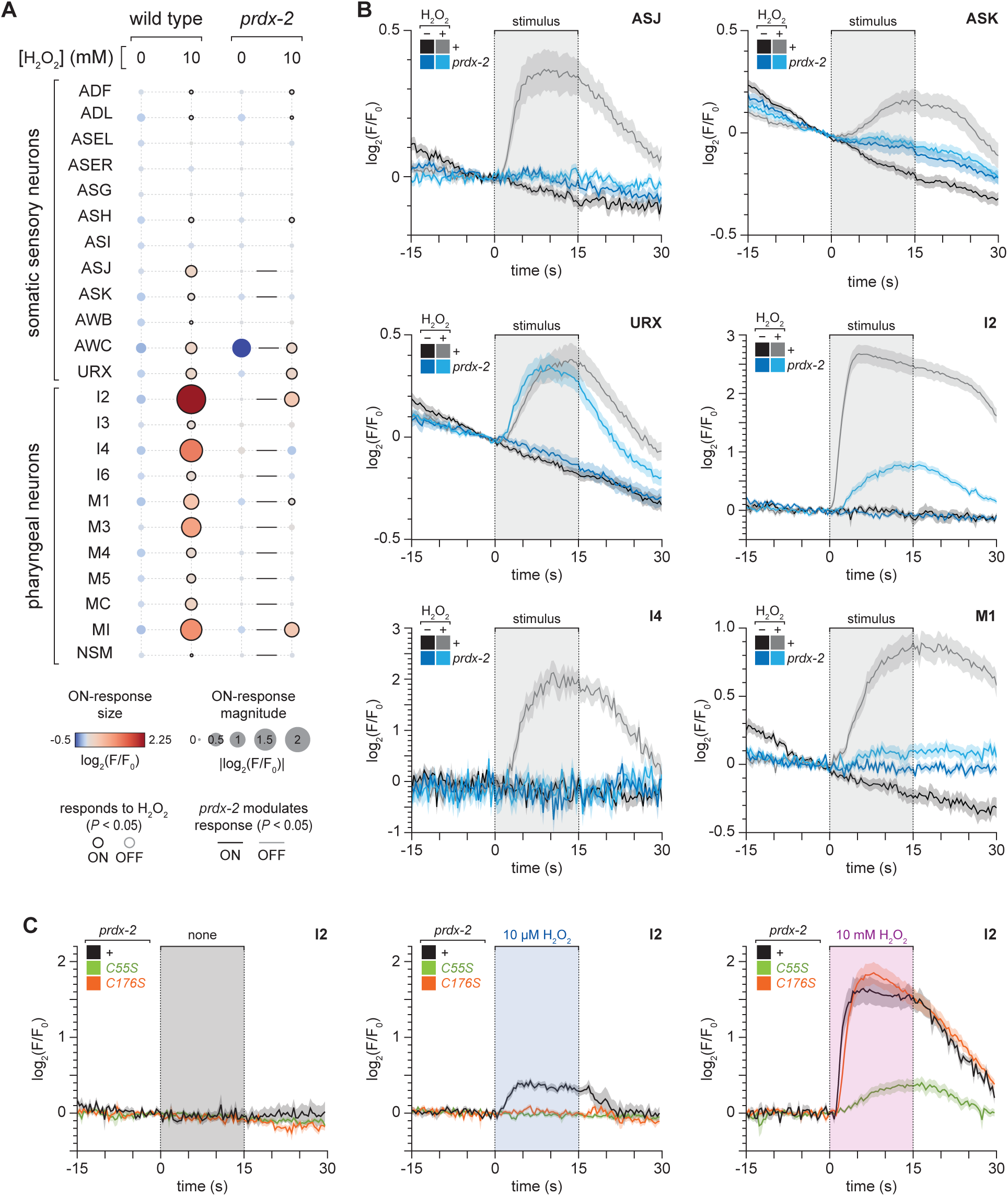
Perception of H_2_O_2_ by sensory neurons in both the somatic and pharyngeal nervous systems requires the cytosolic peroxiredoxin PRDX-2. (A) Responses of somatic and pharyngeal neuron classes to 0 and 10 mM H_2_O_2_ in wild-type animals and *prdx-2(gk169)* mutants. Each circle represents one neuron class at a given H_2_O_2_ concentration. Circle color encodes the mean change in GCaMP6 fluorescence during the 15-second stimulus interval (ON response); circle area scales with the absolute response magnitude. Black rings indicate ON responses that differ significantly from the no-H_2_O_2_ control (*t*-test, *P* < 0.05); gray rings indicate significant OFF-responses measured during the 15-second post-stimulus interval (*t*-test, *P* < 0.05); unmarked circles indicate responses that were not significant in either interval (*P* > 0.05). Black and grey lines indicate that ON– and OFF-responses, respectively, are affected by *prdx-2* (standard least-squares regression, *P* < 0.05), otherwise not significant (*P* > 0.05). (B) Mean GCaMP6 fluorescence traces for ASJ, ASK, URX, I2, I4, and M1 neuronal classes in response to 0 and 10 mM H_2_O_2_ in wild-type animals and *prdx-2(gk169)* mutants. Data are plotted as mean ± s.e.m. The shaded box marks the stimulus delivery interval. The remaining neuronal classes are shown in Figure S5. (C) Mean GCaMP6 fluorescence traces for I2 neurons in response to 0, 10 µM, and 10 mM H_2_O_2_ in wild-type animals, and in *prdx-2(C55S)* and *prdx-2(C176S)* mutants. Data are plotted as mean ± s.e.m. The shaded box marks the stimulus delivery interval. Statistical analysis for panels (A-B) is in Table S7, and for panel (C) in Table S8.

We next asked how PRDX-2’s catalytic cysteines contribute to neuronal H_2_O_2_ sensing. We focused on the I2 neurons because they are the most H_2_O_2_-sensitive neurons. Because C55 is the first residue to react with H_2_O_2_ in the PRDX-2 catalytic cycle, we expected it to be essential for H_2_O_2_-evoked activity. For C176, the resolving cysteine, several outcomes were possible: it could be required at all H_2_O_2_ concentrations if C_P_-C_R_ disulfide formation is needed for signaling; required only at low H_2_O_2_, where a long-lived C_P_-C_R_ intermediate might support sensing when sulfenylated C55 is rare; largely dispensable if sulfenylated C55 alone suffices; or, conversely, its loss could enhance signaling by increasing the half-life of sulfenylated C55. To test the roles of these residues, we measured I2 calcium responses to 0, 10 µM, and 10 mM H_2_O_2_ in wild-type worms and in *prdx-2(C55S)* and *prdx-2(C176S)* mutants. Both mutations abolished the response to 10 µM H_2_O_2_; however, at 10 mM H_2_O_2_, C55S sharply reduced the response but C176S had no effect (Figure 5C). Thus, even though both of PRDX-2’s catalytic cysteines are necessary for chemotactic avoidance of H_2_O_2_, they contribute to neuronal H_2_O_2_ sensing in distinct ways. The peroxidatic cysteine was indispensable for PRDX-2-mediated H_2_O_2_ perception in I2, while the resolving cysteine was only necessary at low H_2_O_2_ concentrations, consistent with a model in which the C_P_-C_R_ disulfide intermediate supports sensing under limiting H_2_O_2_.

During these experiments, we also unexpectedly discovered that the C-terminus of PRDX-2 plays a role in H_2_O_2_ perception and avoidance. We could not restore H_2_O_2_ avoidance in *prdx-2(−)* mutants expressing a translational reporter, *nEx2078[prdx-2p::prdx-2(+)-mCherry]*, which consists of a C-terminal fusion of mCherry to a *prdx-2* genomic fragment, including its promoter and coding sequence^22^ (Figure S6A), suggesting that this visibly expressed fusion protein^22^ was inactive. We hypothesized that this lack of rescue resulted from a dominant-negative effect, since previous studies showed the C-terminus can be crucial in the 2-Cys PRX catalytic cycle: when 2-Cys PRXs are in a reduced form, their C-terminus binds to residues on the opposing subunit, stabilizing the assembly of these dimers into decamers^88^ and slowing disulfide bond formation^72^; but upon sulfenylation of C_P_ by H_2_O_2_, the C-terminus becomes unstructured, allowing specific proteins to bind it^89^ and also triggering the dissociation of decamers into disulfide-bonded dimers^72^. Supporting this idea, expressing *nEx2078* in a wild-type background lowered H_2_O_2_ avoidance (Figure S6A), an effect not observed in *prdx-2(−)* mutants or by overexpressing mCherry alone from the *eft-3* promoter (Figure S6A). Moreover, we examined the activity of somatic sensory neurons in response to 10 mM H_2_O_2_ in animals expressing *nEx2078* in a wild-type background and found reduced or absent GCaMP responses in ASH, ASK, and ASJ neurons (Figure S6B-C, Figure S7). We conclude that the PRDX-2 C-terminus is important for PRDX-2 function and that its fusion to mCherry dominantly interferes with wild-type PRDX-2 activity required for H_2_O_2_ perception and avoidance.

We showed that many sensory neurons, both pharyngeal and somatic, required the cytosolic enzyme PRDX-2 to increase intracellular calcium when extracellular H_2_O_2_ increased. In wild-type animals, increasing extracellular H_2_O_2_ elicited progressively larger calcium signals in I2 and other H_2_O_2_-sensing neurons. If PRDX-2 merely degraded H_2_O_2_, its loss should cause higher intracellular H_2_O_2_ levels and therefore larger neuronal responses. Instead, *prdx-2(−)* mutants showed markedly reduced or absent calcium responses to H_2_O_2_. Given that PRXs are abundant and their peroxidatic cysteine is seven orders of magnitude more reactive toward H_2_O_2_ than typical solvent-exposed cysteines^90^, it is unlikely that H_2_O_2_ directly oxidizes target proteins. We therefore propose that PRDX-2 functions as an active transducer of the H_2_O_2_ signal, such that oxidation of its peroxidatic (C55) and resolving (C176) cysteines by H_2_O_2_ conveys information to downstream effectors that trigger the calcium response (Figure 4D).

### Chemotactic avoidance of H_2_O_2_ is driven by somatic and pharyngeal neuronal sensing through the ion channels LITE-1 and GUR-3

Because PRDX-2 is a cytosolic enzyme, its oxidation by H_2_O_2_ must be communicated to membrane ion channels that drive neuronal activation and avoidance. To identify which proteins mediate the PRDX-2-dependent response in somatic and pharyngeal neurons, we used a candidate-gene approach focused on the light-activated ion channels LITE-1 and GUR-3^22,26,91,92^, members of the insect gustatory receptor family that assemble as tetrameric ion channels^26,93^. LITE-1 and GUR-3 mediate aversive responses—including body reversals, feeding inhibition, and spitting—in response to UV/blue light and H_2_O_2_ ^22,24,91,92^. *lite-1(ce314)* null mutants and *gur-3(ok2245)* null mutants lowered the H_2_O_2_ avoidance index by 0.16 and 0.32, respectively, compared to wild-type controls (Figure 6A). *lite-1(ce314) gur-3(ok2245)* double mutants avoided H_2_O_2_ to the same extent as *gur-3* single mutants but retained a substantial ability to avoid H_2_O_2_ (Figure 6A). We conclude that LITE-1, GUR-3, and additional mechanisms contribute to H_2_O_2_ avoidance.

**Figure 6.**
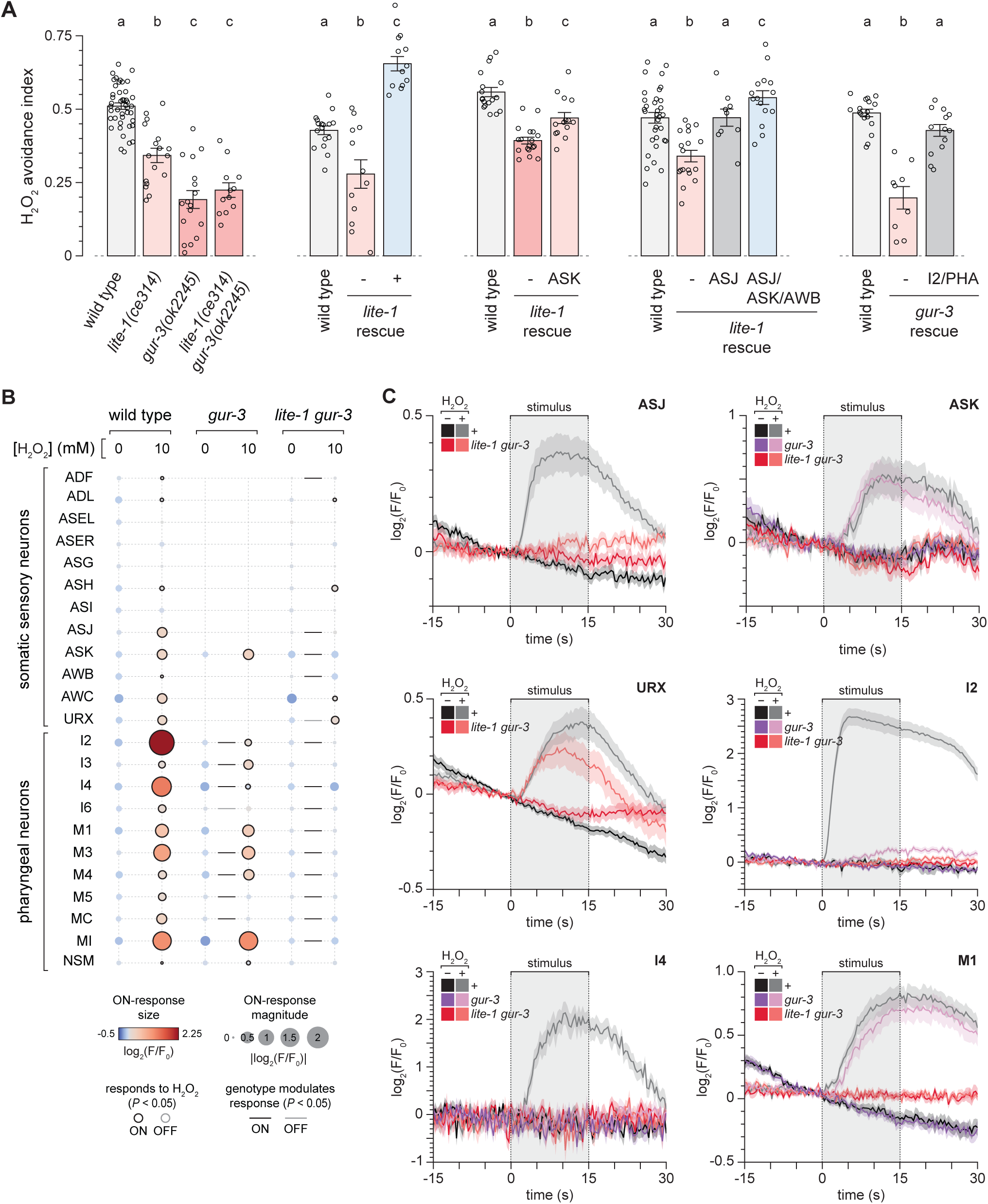
Chemotactic avoidance of H_2_O_2_ is driven by somatic and pharyngeal neuronal sensing through the ion channels LITE-1 and GUR-3. (A) H_2_O_2_ avoidance was lowered by mutations in *lite-1* and *gur-3*. Transgenic rescue experiments of these two mutants using different neuron-class specific promoters. Data are plotted as mean ± s.e.m. Groups labeled with different letters exhibited significant differences (*P* < 0.05, Tukey HSD test), otherwise not significant (*P* > 0.05). (B) Responses of somatic and pharyngeal neuron classes to 0 and 10 mM H_2_O_2_ in wild type animals, in *gur-3(ok2245)* mutants, and in *lite-1(ce314) gur-3(ok2245)* double mutants. Each circle represents one neuron class at a given H_2_O_2_ concentration. Circle color encodes the mean change in GCaMP6 fluorescence during the 15 second stimulus interval (ON response); circle area scales with the absolute response magnitude. Black rings indicate ON responses that differ significantly from the no-H_2_O_2_ control (*t*-test, *P* < 0.05); gray rings indicate significant OFF-responses measured during the 15 second post-stimulus interval (*t*-test, *P* < 0.05); unmarked circles indicate responses that were not significant in either interval (*P* > 0.05). Black and grey lines indicate that ON– and OFF-responses, respectively, are affected by the specific genotype relative to wild type (standard least-squares regression, *P* < 0.05), otherwise not significant (*P* > 0.05). (C) Mean GCaMP6 fluorescence traces for ASJ, ASK, URX, I2, I4, and M1 neuronal classes in response to 0 and 10 mM H_2_O_2_ in wild type animals, in *gur-3(ok2245)* mutants, and in *lite-1(ce314) gur-3(ok2245)* double mutants. Data are plotted as mean ± s.e.m. The shaded box marks the stimulus delivery interval. The remaining neuronal classes are shown in Figures S8. Statistical analysis is in Table S10.

Reintroducing the *lite-1(+)* gene into *lite-1(ce314)* mutants increased H_2_O_2_ avoidance beyond wild-type levels, indicating that LITE-1 promoted H_2_O_2_ avoidance in a dose-dependent manner (Figure 6A). LITE-1 is expressed at least 44 cell types^22,87,92^, including the ASK and ASJ neurons where it mediates phototransduction^94^. Consistent with this expression pattern, restoring *lite-1(+)* expression in ASJ alone, ASK alone, or jointly in ASJ, ASK, and AWB neurons fully rescued H_2_O_2_ avoidance in *lite-1(xu7 R401H)* mutants (Figure 6A). GUR-3 is expressed in the pharyngeal neurons I2, I4, M1, MI, and the interneuron AVD^22,24^; expression of *gur-3(+)* exclusively in I2 and PHA neurons restored H_2_O_2_ avoidance to *gur-3(ok2245)* mutants (Figure 6A). We conclude that hydrogen peroxide avoidance is mediated, at least in part, by LITE-1 in the somatic sensory neurons ASK and ASJ, and by GUR-3 in the pharyngeal neuron I2.

To determine whether *lite-1* and *gur-3* mediate H_2_O_2_ perception by these sensory neurons, we analyzed neuronal responses to 10 mM H_2_O_2_ in mutants lacking one or both genes, as part of the same experiment assessing the requirement of *prdx-2*. Specifically, we investigated in *gur-3(−)* single mutants the responses of H_2_O_2_-sensing pharyngeal neurons and the ASK neuron (which might express very low levels of GUR-3 based on single-cell mRNA sequencing^87^), and examined in *lite-1(ce314) gur-3(ok2245)* double mutants the responses of both somatic and pharyngeal H_2_O_2_-sensing neurons. The *gur-3* mutation reduced the I2 response by 4.8-fold without fully eliminating it, abolished or nearly abolished the responses of I4, M5, and MC, decreased the M3 response by half, and did not lower the responses of ASK, I3, M1, M4, and MI (Figure 6B, Figure S8). In contrast, the combined loss of *lite-1* and *gur-3* also eliminated the responses of ADF, ASJ, ASK, I2, I3, M1, M3, M4, and MI (Figure 6B, Figure S8). Loss of both these genes did not affect responses in ADL, ASH, and URX (Figure 6B, Figure S8). Thus, the combined functions of GUR-3 and LITE-1 were essential for H_2_O_2_ perception by pharyngeal (I2, M1) and somatic (ASK and ASJ) neurons that promoted H_2_O_2_ avoidance.

### GUR-3 cysteines mediate H_2_O_2_ perception by I2 neurons leading to chemotactic avoidance

Finally, we investigated how GUR-3 ion channels sensed H_2_O_2_ to drive chemotactic avoidance behavior. Previous structural modeling showed that LITE-1 and GUR-3 share highly similar homo-tetrameric structures^26^. Structure-function analyses of LITE-1 suggested two mechanisms for its response to light: direct sensing through a chromophore bound to cysteine 300, which faces the channel’s pore^26^, and indirect sensing through oxidation of cysteine 258, a cytosol-facing residue that is thought to form a disulfide bond with peroxiredoxin in response to photochemically generated cytosolic H_2_O_2_^22,26,76^. We used CRISPR gene editing to mutate the corresponding cysteines in GUR-3 to serines (Figure 7A). The *gur-3(C260S)* mutation, which affected the putative PRDX-2 binding cysteine, reduced H_2_O_2_ avoidance to the same extent as the null allele of the gene (Figure 7A). Similarly, the *gur-3(C301S C302S)* mutation, affecting the two putative chromophore-binding cysteines, lowered the H_2_O_2_ avoidance index by 0.22 (Figure 7A); this reduction was not observed in *gur-3(C301S C302S)/+* heterozygotes (Figure 7A), indicating that the mutation was recessive. We conclude that both the putative chromophore-binding and PRDX-2-binding cysteines in GUR-3 promoted H_2_O_2_ avoidance.

**Figure 7.**
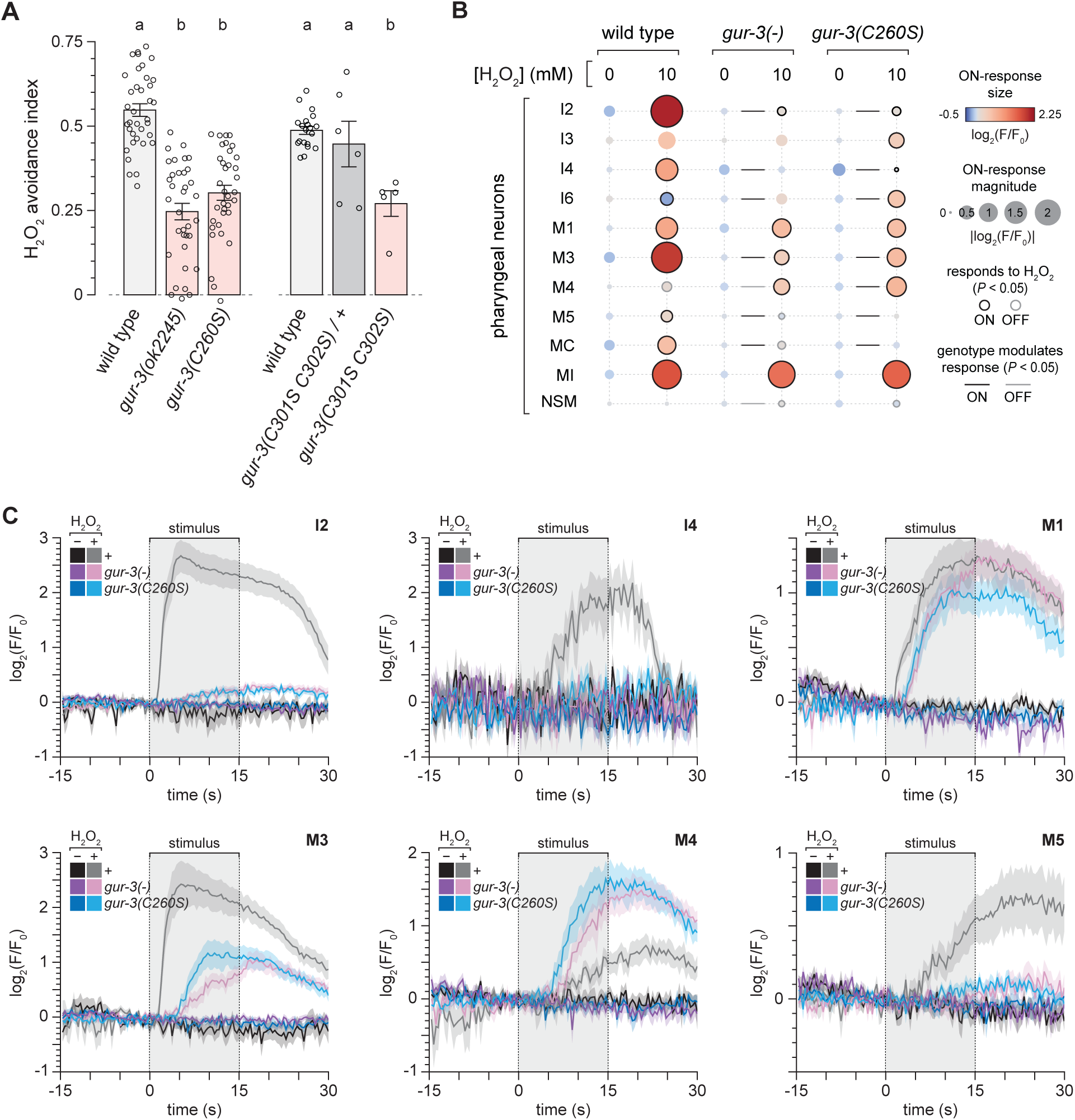
GUR-3 cysteines mediate H_2_O_2_ perception by I2 neurons leading to chemotactic avoidance. (A) Replacing the C260 residue of *gur-3* with serine lowered H_2_O_2_ avoidance to *gur-3* null levels, while replacing the C301 and C302 residues with serine caused a recessive decrease in avoidance. Data are plotted as mean ± s.e.m. Groups labeled with different letters exhibited significant differences (*P* < 0.05, Tukey HSD test), otherwise not significant (*P* > 0.05). (B) Responses of somatic and pharyngeal neuron classes to 0 and 10 mM H_2_O_2_ in wild type animals, in *gur-3(ok2245)* mutants, and in *gur-3(C260S)* mutants. Each circle represents one neuron class at a given H_2_O_2_ concentration. Circle color encodes the mean change in GCaMP6 fluorescence during the 15 second stimulus interval (ON response); circle area scales with the absolute response magnitude. Black rings indicate ON responses that differ significantly from the no-H_2_O_2_ control (*t*-test, *P* < 0.05); gray rings indicate significant OFF-responses measured during the 15 second post-stimulus interval (*t*-test, *P* < 0.05); unmarked circles indicate responses that were not significant in either interval (*P* > 0.05). Black and grey lines indicate that ON– and OFF-responses, respectively, are affected by the specific genotype relative to wild type (standard least-squares regression, *P* < 0.05), otherwise not significant (*P* > 0.05). (C) Mean GCaMP6 fluorescence traces for I2, I4, M1, M3, M4, and M5 neuronal classes in response to 0 and 10 mM H_2_O_2_ in wild type animals, in *gur-3(ok2245)* mutants, and in *gur-3(C260S)* mutants. Data are plotted as mean ± s.e.m. The shaded box marks the stimulus delivery interval. The remaining neuronal classes are shown in Figures S9. Statistical analysis is in Table S11.

We also determined the extent to which the thiol group of cysteine 260 in GUR-3 was necessary for GUR-3-dependent responses of pharyngeal neurons to 10 mM H_2_O_2_. The *gur-3(C260S)* mutation reduced I2’s response by 4.5-fold, identical to the reduction caused by the *gur-3* null allele (Figure 7B-C), and mimicked the effect of the *gur-3* null allele in all other pharyngeal neurons (Figure 7B-C, Figure S9). We conclude that the redox-active thiol of cysteine 260 plays an essential role in GUR-3’s ability to affect the activity of pharyngeal neurons in response to H_2_O_2_.

We propose that in response to increases in cytosolic H_2_O_2_, PRDX-2 directly activates the GUR-3 and LITE-1 ion channels via oxidative modification of their cytosol-facing cysteine thiols. Thus, neuronal activity is triggered by the flow of oxidative equivalents from H_2_O_2_ to the channel’s cysteine thiols through an electron relay involving PRX’s C_P_ and C_R_ cysteines.

### Glutamate from pharyngeal and somatic sensory neurons promotes avoidance of H_2_O_2_

To understand how H_2_O_2_-sensing neurons in the pharyngeal and somatic nervous systems directed the animal to navigate away from H_2_O_2_, we examined the types of neurotransmitters they use to communicate with other cells. We focused on glutamate because 13/24 classes of H_2_O_2_-sensing neurons use that neurotransmitter^95^ (Table S5). Glutamate is loaded into synaptic vesicles by the EAT-4 transporter^96^. We found that both *eat-4(ky5)* null mutants and *eat-4(nj2)* loss-of-function mutants showed a 0.4 reduction in H_2_O_2_ avoidance index compared to wild-type controls (Figure 8A). Several neurons that mediated H_2_O_2_ escape are cholinergic (ASJ, AWB, IL2, I1, M1), but *unc-17* mutants, which lack the vesicular acetylcholine transporter^97^, are so uncoordinated that they could not be tested in our chemotactic avoidance assay. Mutants defective in the biosynthesis of other neurotransmitters—including *tph-1(n4622)* for serotonin^98,99^, *cat-2(e1112)* for dopamine^100^, *tdc-1(ok914)* for tyramine^101^, and *tbh-1(ok1196)* for octopamine^101^—did not affect H_2_O_2_ avoidance, but GABA biosynthesis defective *unc-25(e156)*^102^ and GABA transport defective *unc-47(e307)*^103^ lowered H_2_O_2_ avoidance significantly (Figure S10A).

**Figure 8.**
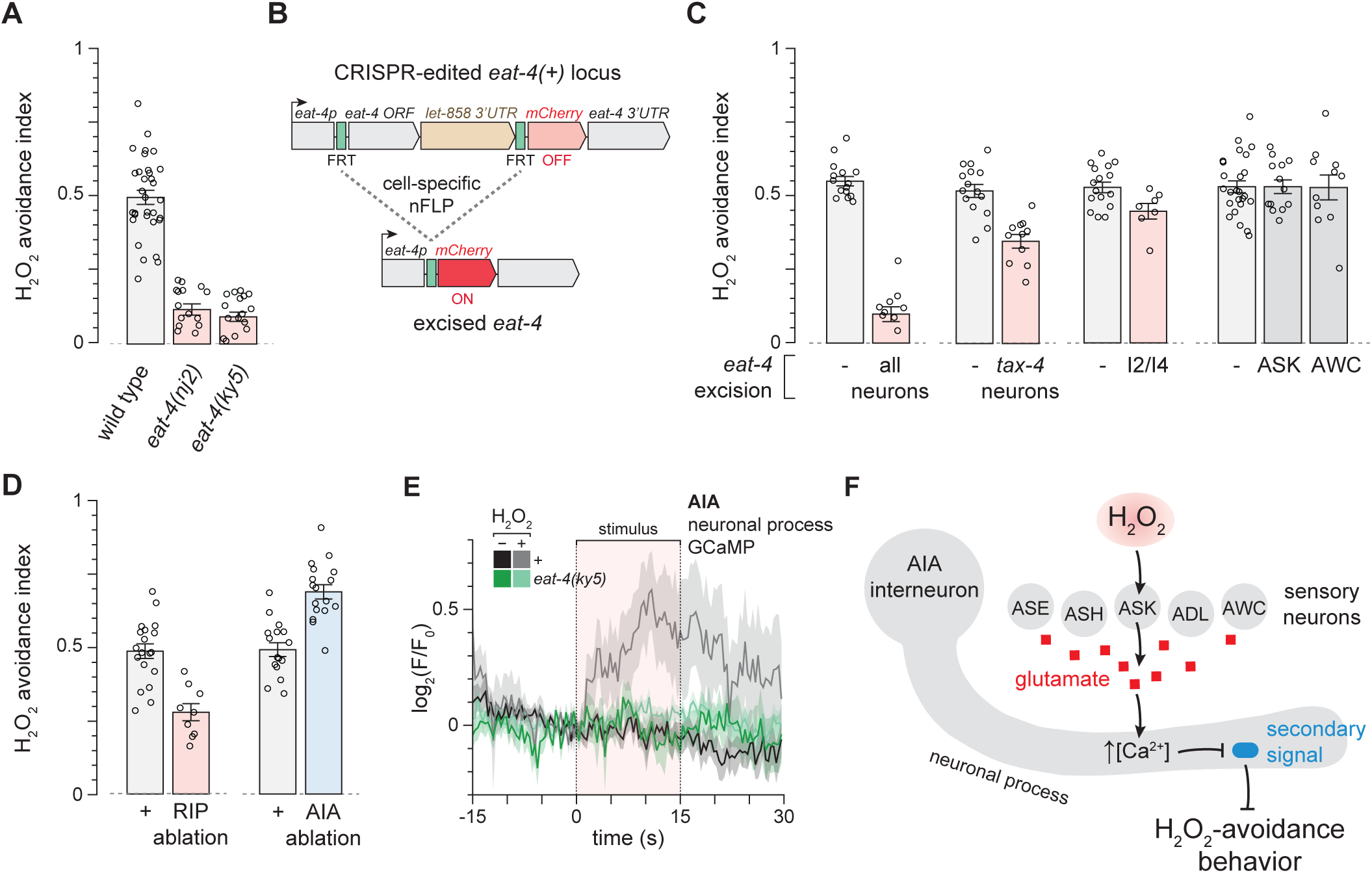
Glutamate from pharyngeal and somatic sensory neurons promotes avoidance of H_2_O_2_. (A) *eat-4* mutants lowered H_2_O_2_ avoidance. (B) Schematic of the cell-selective glutamate-knockout strategy. FRT sites flank the *eat-4* locus; nFLP recombinase expression in specified cells excises the *eat-4* open reading frame (ORF). UTR, untranslated region; FRT, nFLP target sequence. (C) H_2_O_2_ avoidance of nematodes with *eat-4* excised in the specified subsets of neurons. (D) Ablation of RIP interneurons lowered H_2_O_2_ avoidance, while ablation of AIA interneurons increased it. (E) Mean GCaMP6 fluorescence traces in the AIA process in response to 0 and 10 mM H_2_O_2_ in wild type animals and *eat-4(ky5)* mutants. Data are plotted as mean ± s.e.m. The red-shaded box marks the stimulus delivery interval. Statistical analysis is in Table S12. (F) Sensory neurons use glutamate to excite the process of the AIA interneurons in response to H_2_O_2_. This excitation suppresses AIA’s output (secondary signal), producing a net increase in H_2_O_2_ avoidance.

To selectively inactivate glutamatergic neurotransmission, we used a CRISPR-edited version of the endogenous *eat-4* locus that enables excision of the *eat-4* coding sequence and expression of the fluorescent protein mCherry in cells that express a nuclear-localized flippase recombinase (nFLP)^104^ (Figure 8B). Knocking out *eat-4* in all neurons lowered the H_2_O_2_ avoidance index by 0.45 (Figure 8C). Restricting the knockout to 10 classes of somatic ciliated sensory neurons—including the glutamatergic neurons AFD, ASE, ASG, ASK, AWC, and BAG—by expressing nFLP with the *tax-4* promoter reduced the avoidance index by 0.17 (Figure 8C). Selective knockout of *eat-4* in the glutamatergic I2 and I4 neurons^95^ was performed with the cGAL system^105^: cGAL driven from the *gur-3* promoter activated a *UAS::nFLP* effector transgene in both I2 and I4; this manipulation lowered the avoidance index by 0.08 (Figure 8C). *eat-4* knockout only in ASK or AWC neurons had no effect (Figure 8C). Expression of *eat-4(+)* only in I2 and PHA neurons did not affect the H_2_O_2_ avoidance index of *eat-4(ky5)* mutants (Figure S10B). We conclude that both pharyngeal (I2 and/or I4) neurons and a subset of the somatic neurons AFD, ASE, ASG, ASK, and AWC, and possibly additional neurons, promoted hydrogen peroxide avoidance by releasing the neurotransmitter glutamate.

To identify interneurons that may regulate H_2_O_2_ avoidance in response to glutamate and other sensory signals, we focused on the RIP interneurons, which form the sole direct synaptic bridge between the somatic and pharyngeal nervous systems^27,40,106^. RIP interneurons receive input from multiple glutamatergic and multiple H_2_O_2_-sensing somatic neurons, connect via gap junctions with the pharyngeal neuron I1 and receive synaptic input from M1 (both of which promoted H_2_O_2_ avoidance). To determine whether RIP was functionally required for H_2_O_2_ avoidance, we ablated the neurons using the cGAL system^105^, driving cGAL from the *nlp-51* promoter in animals carrying a *UAS::caspase* effector transgene. RIP ablation lowered the H_2_O_2_ avoidance index by 0.2 (Figure 8D), indicating that this interneuron promotes chemotactic avoidance of H_2_O_2_. We also asked whether H_2_O_2_ modulated RIP activity. The nucleus of the RIP neurons showed no detectable calcium response to 10 mM H_2_O_2_ in our earlier pan-neuronal GCaMP recordings (Figure S10C), though it remains possible that RIP’s neurite may respond to the stimulus. We propose that RIP-mediated communication between the somatic and pharyngeal nervous systems facilitates H_2_O_2_-avoidance behavior in *C. elegans*.

In parallel, we investigated the AIA interneurons, which participate in chemotaxis and foraging^45,104,107,108^ and receive extensive glutamatergic input^40,47,95^. Glutamate released from ADL, ASE, ASH, ASK, and AWC can influence AIA activity likely through glutamate-gated chloride channels^45,104,108,109^. We found that AIA ablation increased H_2_O_2_ avoidance index by 0.2 (Figure 8D), indicating that this interneuron class normally inhibits chemotactic avoidance of H_2_O_2_. To determine whether AIA activity is modulated by H_2_O_2_, we measured this interneuron’s activity using the same microfluidic setup as before, but this time using nematodes expressing GCaMP6s exclusively in AIA. This approach allowed us to record AIA activity specifically in the neurite (rather than the nucleus), as calcium transients in AIA and other interneurons are predominantly localized to the neuronal process due to its high density of synaptic connections^107,110–112^. Exposure to 10 mM H_2_O_2_ increased AIA activity in the process, and these effects were absent in *eat-4(ky5)* mutants (Figure 8E). Therefore, environmental H_2_O_2_ excites AIA’s process through a mechanism that requires vesicular loading of glutamate by the EAT-4 transporter.

We propose that upon modulation by H_2_O_2_, many classes of sensory neurons signal via glutamate to excite AIA (Figure 8F). Because ablating AIA increases H_2_O_2_ avoidance, we infer that AIA sends a secondary signal that inhibits H_2_O_2_ avoidance, and that glutamate-dependent AIA excitation represses this inhibitory output, resulting in a net pro-avoidance effect (Figure 8F).

## Discussion

Hydrogen peroxide (H_2_O_2_) is the most common reactive chemical threat faced by organisms. Here, we mapped the neural circuit that drives chemotactic escape from environmental H_2_O_2_ in *C. elegans* and identified molecular steps that convert changes in intracellular peroxide handling into neural activity and locomotory output. Together, our findings support a model in which the nervous system functions as a hydrogen peroxide sentinel: a distributed set of sensory neurons in the mouth and nose monitors the state of the cytosolic H_2_O_2_-detoxification machinery and relays this information to interneurons that drive escape behavior. An important implication is that peroxide detection and peroxide detoxification are mechanistically coupled. Whether similar coupling underlies neural responses to other reactive chemicals is an open question.

### Distributed H_2_O_2_ sensors with tiered detection thresholds build a robust chemotactic escape circuit

Our results show that *C. elegans* uses a large sensory array to monitor H_2_O_2_. We identified 24 neuron classes excited or inhibited by environmental H_2_O_2_, extending previous findings by our lab and others^21,22,25^. Twelve of these classes reside in the amphid and labial organs projecting to the nose, where their ciliated endings contact the exterior of the animal or the pseudocoelomic body cavity. Twelve classes are pharyngeal; their non-ciliated endings span successive segments of this organ—from the mouth opening through the corpus, sieve, isthmus, grinder, and terminal bulb—either contacting the lumen or embedded in deeper tissue layers. In addition, PHA neurons in the tail also detect H_2_O_2_ ^25^. Together, this distributed array detects H_2_O_2_ that reaches the head or tail, accompanies ingested bacteria, or diffuses into the body (Figure 3L).

This array of sensors drives navigation in spatial gradients of H_2_O_2_. Our targeted ablations showed that seven H_2_O_2_-sensing neuronal classes, five projecting to the nose (AFD, ASH, ASJ, ASK, AWB) and two projecting to the mouth (I2, M1), each promote escape; and IL2 and I1 also contribute despite showing no calcium responses in our assay. Importantly, eliminating any single one of these classes reduced—but never abolished—avoidance. Thus, the circuit is robust to the loss of individual inputs, consistent with a redundant sensory architecture in which overlapping detectors limit performance degradation when one input fails^113^.

The H_2_O_2_-detection array also partitions H_2_O_2_ concentrations into risk tiers. I2 at the mouth is the most sensitive, responding at 1 µM H_2_O_2,_ consistent with an early warning role. ASJ, M4, and MI engage at 100 µM, possibly relaying rising danger at a dose that remains sublethal. Higher-threshold sensors such as ASK and M1 activate at 1 mM, a level that arrests larval development and shortens lifespan tenfold^17–19,21,32,114^. Additional tiers respond at 3, 10, and 100 mM, lethal on progressively shorter timescales. Several neuronal classes encode exposure history by desensitizing or becoming refractory to H_2_O_2_, so their activity reflects both instantaneous concentration and recent dynamics. These adaptive mechanisms likely prevent saturation during sudden surges: as low-threshold sensors adapt, high-threshold sensors take over, preserving H_2_O_2_-gradient information. We propose that the anatomical arrangement and tiered sensitivities of this array allow animals to measure both how much peroxide is present and how much is about to be ingested.

Combining this and prior work^21,22,25^, we and others evaluated 30 sensory-neuron classes for H_2_O_2_-evoked activity. Within this set, nine of ten amphid, labial, and phasmid classes that drive locomotory avoidance when excited by chemical, mechanical, or thermal cues were also excited by H_2_O_2_, while one class that promotes attraction when excited was inhibited by H_2_O_2_ (Table S5). H_2_O_2_ also excited four of the five pharyngeal neurons mediating UV light-induced feeding inhibition and spitting^22–24^ (Table S5). Consistent with a broader pattern, in *Drosophila* UV light elevates H_2_O_2_, which excites antennal olfactory neurons and suppresses oviposition^115^. Together, these observations suggest that H_2_O_2_-evoked excitation is the prevailing response among aversion-driving sensory neurons. This prevalence could reflect an ancestral H_2_O_2_-sensing modality retained during diversification, or repeated co-option of H_2_O_2_ as a general danger co-signal that yields adaptive value through a shared threat computation. It will be interesting to distinguish between these alternatives by mapping the sensory-neuronal responses to H_2_O_2_ across taxa.

### Multiple communication channels from sensors in the mouth and nose relay information about H_2_O_2_ levels to promote escape

The H_2_O_2_-detection array drives chemotactic escape by relaying information through multiple channels. Glutamate released from subsets of pharyngeal and amphid neurons drives H_2_O_2_ avoidance, with each set making a distinct, nonredundant contribution. In addition, other pharyngeal and amphid neurons in the array are non-glutamatergic and likely signal via acetylcholine, other secreted signals, or electrical coupling (Table S5). This mixed strategy provides transmission fault tolerance: if one signal fails, another still carries the message.

To determine how these signals modulate H_2_O_2_ chemotactic escape, we used a candidate approach and identified two interneuron classes with opposing roles: AIA prevents escape, while RIP promotes escape. We found that H_2_O_2_ excites AIA through a mechanism that requires glutamate release. AIA operates as a coincidence detector: it does not respond reliably to a single sensory input but instead changes state when coordinated inputs arrive^108^. We propose that, in animals exposed to H_2_O_2_, coordinated glutamatergic input from AIA’s presynaptic partners^40,47,95^, including H_2_O_2_-sensing ASH and ASK, excites AIA via glutamate to promote avoidance (Figure 8F). Additional interneuron classes such as AIB, AIY, AIZ, RIM, and RMG, form chemical synapses and gap junctions with H_2_O_2_-sensing amphid neurons that promote escape^40,47,95^ and may translate these inputs into chemotactic avoidance.

While neurons projecting to the nose are well known to drive chemotaxis to attractive and repulsive cues^28^, no pharyngeal neuron class was previously implicated in chemotaxis. We found that the I1, I2, and M1 neuronal classes in the pharynx promote H_2_O_2_ chemotactic avoidance, with glutamate released from I2 and I4 contributing to the behavior. Glutamate is thought to act over short distances and not diffuse beyond the synapse, except under conditions that promote spillover due to impaired glutamate reuptake by adjacent cells^116,117^. How does glutamate from I2, and signals from I1 and M1, modulate the somatic interneurons that drive locomotion? Notably RIP, the sole bridge between the pharyngeal and somatic nervous systems^27,40,106^, promotes H_2_O_2_ avoidance. The two RIP interneurons are well positioned via gap-junction coupling to relay somatic inputs to shape the output of I1; whether they also transmit signals in the opposite, pharyngeal-to-somatic direction remains unknown.

Previous studies show that two pharyngeal circuits, I2 with I4 and M1, and RIP with I1 and MC, respond to UV light and H_2_O_2_ by inhibiting pumping and promoting spitting^22–24^. These findings suggest an additional route by which H_2_O_2_ perception by I2, I4, M1, and MC could influence whole-body escape: proprioceptive feedback. In a pressurized hydrostat like *C. elegans*, pharyngeal contractions could generate pressure waves that spread through the body and be sensed by mechanosensitive cells that modulate the locomotory circuit. This would be similar to findings that motor neuron pressure sensing drives forward locomotion^118^ and influences the state of interneurons driving locomotion^119^.

### Neuronal perception of environmental H_2_O_2_ is controlled by the internal state of the cytosolic machinery that degrades H_2_O_2_

To understand the mechanisms by which the neuronal H_2_O_2_-detection array senses H_2_O_2_ to drive chemotactic escape, we used a combination of molecular genetics, neuronal calcium imaging, and behavioral assays. This approach identified proteins acting in defined neuron classes and pinpointed specific cysteine residues required for their function in H_2_O_2_ perception and avoidance, leading us to propose the following model: The increase in cytosolic H_2_O_2_ following environmental exposure rapidly oxidizes the peroxidatic cysteine of PRDX-2 (C55) to a sulfenic acid (–SOH) intermediate; from there, this intermediate can follow three fates: transmission, recycling, and inactivation (Figure 9).

**Figure 9.**
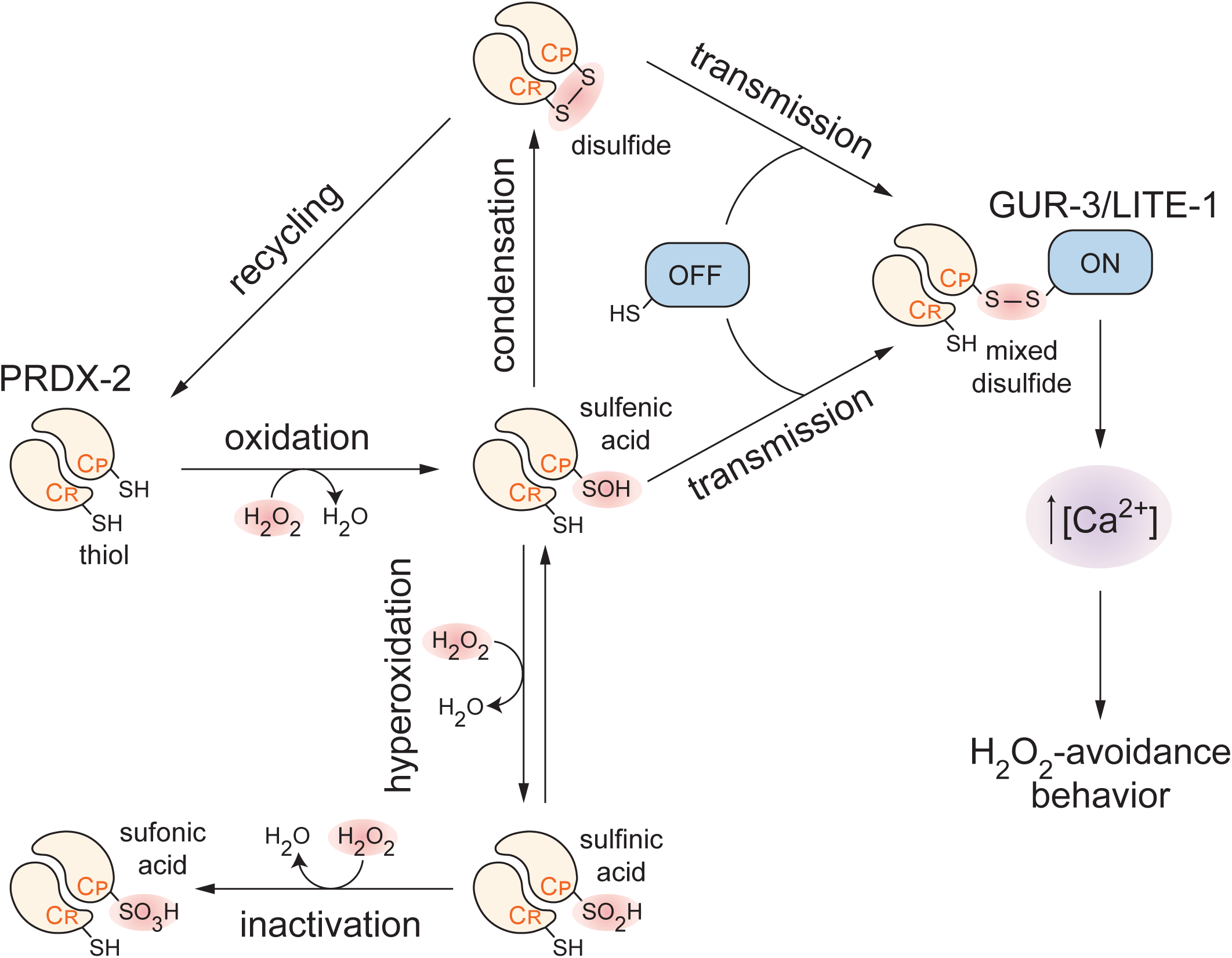
Neuronal perception of environmental H_2_O_2_ is controlled by the internal state of the cytosolic machinery that degrades H_2_O_2_. Environmental H_2_O_2_ oxidizes the peroxidatic cysteine (C_P_) of the cytosolic peroxiredoxin PRDX-2 in specific H_2_O_2_-sensing neurons. The resulting sulfenylated C_P_ can (i) transmit its oxidative equivalents to specific cytosol-facing cysteines in GUR-3 and LITE-1, opening these channels, raising [Ca^2+^], and triggering H_2_O_2_ avoidance behavior; (ii) be recycled by the thioredoxin system, resetting the PRDX-2 sensor and preserving its transmission capacity by preventing its hyperoxidation by additional H_2_O_2_ molecules; or (iii) undergo irreversible inactivation by hyperoxidation. Additionally, in the first step of recycling, the condensation of sulfenylated C_P_ with the resolving cysteine (C_R_) forms a C_P_-C_R_ disulfide within the PRDX-2 dimer that can relay oxidative equivalents to GUR-3 and LITE-1 channel cysteines via thiol-disulfide exchange.

**Transmission**: the sulfenylated C_P_ of PRDX-2 relays oxidative equivalents to cytosol-facing cysteines on the H_2_O_2_-sensing ion channels GUR-3 and LITE-1 via a condensation reaction, forming disulfide bonds at GUR-3 C260 and LITE-1 C258; this shifts ion channel gating, triggering calcium influx, which promotes escape via downstream effectors. AlphaFold multimer modeling positions PRDX-2 next to cytosolic loops of both LITE-1 and GUR-3, bringing its peroxidatic C55 into contact with LITE-1 C258 and GUR-3 C260, respectively^26^. Because direct biochemical evidence for these contacts is lacking, additional proteins or alternative carriers may relay oxidative equivalents from PRDX-2’s sulfenylated C_P_ to these ion channels’ cysteines. The PRDX-2 resolving cysteine C176 is required for transmission in I2 neurons at low H_2_O_2_ (10 µM) but is dispensable at high H_2_O_2_ (10 mM). We propose that the C_P_-C_R_ disulfide-bonded PRDX-2 dimers formed in the first recycling step constitute a long-lived intermediate that engages in thiol-disulfide exchange with GUR-3 C260 and LITE-1 C258 at low H_2_O_2_. At high H_2_O_2_, sulfenylated C_P_ is sufficiently more abundant to rapidly drive ion channel modification without forming a C_P_-C_R_ intermediate, consistent with proteomic studies in yeast and human cells showing C_P_ is essential and C_R_ modulates the formation of PRX disulfides with many proteins^120,121^.

**Recycling**: the sulfenylated C_P_ of PRDX-2 condenses with C_R_ to form a C_P_-C_R_ disulfide-bonded dimer. The thioredoxin (TRX)-thioredoxin reductase (TRXR-1) redox relay^79^, powered by NADPH, reduces this disulfide, restoring C55 and C176 to thiols and resetting the sensor. Different H_2_O_2_-sensing neuronal classes likely recruit distinct TRX-family members to reset PRDX-2; in ASJ, TRX-1 provides the neuron-class specific activity and operates through its C38 (catalytic) and C72 (non-catalytic). Although recycling lowers the pool of sulfenylated C_P_, it sustains transmission capacity by preventing C_P_ hyperoxidation that would otherwise trap PRDX-2 in an inactive state and by regenerating sensing-competent C_P_.

**Inactivation**: the sulfenylated C_P_ of PRDX-2 can react with H_2_O_2_, yielding hyperoxidized sulfinic (–SO_2_H) and, with sustained exposure, sulfonic (–SO_3_H) acids that inactivate the enzyme and sever the redox relay to LITE-1 and GUR-3. Consistent with this, biochemical studies show that after *C. elegans* nematodes are exposed to 5 mM H_2_O_2_ for 5 minutes, reduced PRDX-2 disappears and is completely replaced by hyperoxidized and disulfide-linked forms; the hyperoxidized PRDX-2 persists for at least 4 hours of recovery, indicating that this state is not readily reversible^69^.

The 2-Cys peroxiredoxin subfamily PRDX-2 alone cannot account for all H_2_O_2_ sensing across the neuronal array. The cytosolic 1-Cys peroxiredoxin subfamily PRDX-6 also promoted H_2_O_2_ avoidance, and we propose that it contributes directly to transmission. Notably, the *gur-3(C260S)* mutation impaired H_2_O_2_ perception more severely than *prdx-2(C55S)*, suggesting that GUR-3 C260 often receives oxidative equivalents from an alternative source rather than exclusively from PRDX-2. We therefore propose that upon oxidation by H_2_O_2_, the sulfenylated C_P_ of PRDX-6 directly relays oxidative equivalents to GUR-3 and LITE-1 cysteines in neuron classes where loss of *prdx-2* did not phenocopy the *lite-1 gur-3* double mutant, including I2, M1, MI, and AWC. Recycling of 1-Cys PRXs depends on the glutathione relay: two GSH molecules reduce the sulfenylated C_P_, and GSR-1 restores GSH from GSSG^73–75^, powered by NADPH; partial loss of *gsr-1* lowered H_2_O_2_ avoidance, supporting a PRDX-6-based transmission route. Moreover, H_2_O_2_ excited URX, ASH, ADL, and AWC via a mechanism independent of PRDX-2, GUR-3, and LITE-1, indicating that additional mechanisms are responsible for H_2_O_2_-dependent excitation in these neurons. We propose that PRDX-2 and PRDX-6 establish parallel redox wires that converge on GUR-3 and LITE-1 ion channel cysteines, providing fault tolerance and preserving transmission when either wire is compromised.

### Peroxiredoxins make it possible for micromolar H_2_O_2_ to modulate neuronal activity

How do *C. elegans* neurons reliably detect H_2_O_2_ concentrations as low as 1 µM on the timescale of seconds? The proteome contains ∼210,000 cysteines^122^ and any of these cysteines could, in principle, sense H_2_O_2_ if oxidation of that residue altered protein function. However, fast and reliable micromolar sensing requires an exceptionally reactive cysteine thiol, because the rate of thiol oxidation is proportional to H_2_O_2_ concentration and falls exponentially with the reaction’s free energy of activation set by each cysteine’s microenvironment^123,124^. We show that *C. elegans* neurons satisfy this kinetic requirement by using one of the proteome’s most reactive cysteines as their H_2_O_2_ sensor: the peroxidatic cysteine in the active site of PRDX-2. The exceptional reactivity of C_P_ towards H_2_O_2_ arises from the electrostatic and hydrogen-bonding environment of the peroxiredoxin active site, which activates both C_P_ and H_2_O_2_, and stabilizes the transition state for their reaction^125^. Because C_P_ reacts with H_2_O_2_ ∼10^7^-fold faster than typical cysteines and ∼10^5^-fold faster than even highly reactive cysteines^126^, peroxiredoxins are the primary enzymes that remove H_2_O_2_ at micromolar concentrations^1^. As a result, C_P_ captures and degrades most H_2_O_2_ molecules before they oxidize less reactive cysteines elsewhere in the proteome^126^. For these reasons, we disfavor models in which low concentrations of H_2_O_2_ directly oxidize cytosol-facing cysteines on LITE-1 or GUR-3 to open these ion channels, with PRDX-2 in its reduced form acting only later to recycle the oxidized cysteines.

Traditionally, H_2_O_2_ has been viewed primarily as a cytotoxic oxidant. At the 0.25-1 mM H_2_O_2_ concentrations reported during inflammation and post-ischemic reperfusion in mammals^127,128^, H_2_O_2_ reacts rapidly with proteins, nucleic acids, and lipids, causing macromolecular damage that leads to neuronal dysfunction and is thought to increase the aging brain’s vulnerability to neurodegenerative disease^129,130^. In contrast, our studies in *C. elegans* show that H_2_O_2_ can quickly modulate neuronal activity at much lower concentrations, through a process enabled by cytosolic peroxiredoxins. Moreover, even transient exposure to H_2_O_2_ can leave a persistent functional effect, since pre-exposure to sub-micromolar H_2_O_2_ potentiates the subsequent glycerol response of ASH in a PRDX-2-dependent manner^131^. This emerging view of peroxiredoxins as signal transducers rather than mere detoxifying enzymes^7^ is also supported by recent proteomic studies in yeast and human cells, which show that peroxiredoxins form evolutionarily conserved disulfides with hundreds of proteins in response to H_2_O_2_^120,121^. Although the full scope of this peroxiredoxin-mediated regulation of the proteome is only beginning to be understood, some interactions are already known to be functionally important^7,120^. Taken together with the facts that production of non-cytotoxic levels of H_2_O_2_ is a commonplace output of cellular metabolism^6^, and that H_2_O_2_ can diffuse over hundreds of microns and cross membranes^6,8^, these observations lead us to propose that peroxiredoxins enable H_2_O_2_ to play a much broader role in regulating neuronal activity than previously appreciated. In this view, cytosolic peroxiredoxins transduce information carried by non-cytotoxic levels of H_2_O_2_ into long-range electrical signals that can modulate behavior and distant physiology. Going forward, it will be interesting to determine whether circuit defects in aging and neurodegenerative disease arise from changes in H_2_O_2_ signaling via peroxiredoxins rather than from direct oxidative damage.

## Materials and Methods

### *C. elegans* culture and strains

*C. elegans* strains were derived from the wild-type strain N2 (Bristol)^132^ and were cultured according to standard laboratory conditions on nematode growth medium (NGM, 17 g/L agar, 2.5 g/L Bacto Peptone, 3.0 g/L NaCl, 1 mM CaCl_2_, 1 mM MgSO_4_, 25 mM H_2_KPO_4_/HK_2_PO_4_ pH 6.0, 5 mg/L cholesterol) seeded with *E. coli* OP50 at 20°C unless noted otherwise. Before experiments, we propagated all strains for at least two generations after recovery from starvation or freezing. Mutant alleles of *gur-3* and *prdx-2* were generated by SunyBiotech using CRISPR/Cas9 genome engineering. The sequences of the wild type and edited alleles are in Supplementary Data 1. We outcrossed mutants and generated double mutants using standard genetic methods. Unless indicated, we obtained chemicals from Sigma. For a list of all worm and bacterial strains used in this study, see Tables S1 and S2, respectively. For a list of PCR genotyping primers and phenotypes used for strain construction, see Table S3.

### Transgenic driver lines using the cGAL expression system

To build the I1 driver line, we constructed pLJ30 [*lgc-8p::NLS::cGAL(DBD)::cGAL(AD)::let-858 3′UTR*] by amplifying *lgc-8p* with primers oLJ117/oLJ118 and cloning the product into pHW393 between the FseI and AscI sites. We injected pLJ30 into PS7149 at 30 ng/µl together with *unc-122p::RFP* (40 ng/µl) to generate INF67 (*syIs390[15xUAS::?pes-10::GFP::let-858 3′UTR + ttx-3p::RFP + 1 kb DNA ladder (NEB)]; nonEx34[lgc-8p::NLS::cGAL(DBD)::cGAL(AD)::let-858 3′UTR + coel::RFP]*). *syIs390* contains a GFP effector that we used to verify the driver line’s expression pattern. We outcrossed *syIs390* from INF67 to obtain INF203 (*nonEx34*), UV-integrated the array, and backcrossed six times to obtain INF125 (*nonIs4[lgc-8p::NLS::cGAL(DBD)::cGAL(AD)::let-858 3′UTR + coel::RFP]*). To build the I2 driver line, we constructed pLJ73 *[aqp-5p::cGAL-N-term]* by amplifying *aqp-5p* with primers oLJ176/oLJ177, cloning the product into pHW393 (FseI/AscI) to generate pLJ33, and subcloning *aqp-5p* into pHW530 (FseI/AscI) to obtain pLJ73. We also constructed pLJ74 *[gur-3p::cGAL-C-term]* by amplifying *gur-3p* with primers oLJ182/oLJ183, cloning the product into pHW393 (FseI/AscI) to generate pLJ34, and subcloning *gur-3p* into pJL081 (FseI/AscI) to obtain pLJ74. We co-injected pLJ73 and pLJ74 into PS7149 at 15 ng/µl each, together with *unc-122p::tagBFP (40 ng/µl)*, to generate INF471 (*syIs390; nonEx150[aqp-5p::cGAL-N, gur-3p::cGAL-C + unc-122p::tagBFP]*).

We then isolated a spontaneous integrant from INF471 and backcrossed it twice to obtain INF499 (*syIs390; nonIs9[aqp-5p::cGAL-N, gur-3p:: cGAL-C + unc-122p::tagBFP]*). To build the RIP driver line, we constructed pLJ51 *[nlp-51p::NLS::cGAL(DBD)::cGAL(AD)::let-858 3′UTR]* by amplifying *nlp-51p* from pLJ47 (unpublished) with primers oLJ275/oLJ262 and cloning the product into pHW393 between the SphI and BamHI sites. We injected pLJ51 into PS7149 at 30 ng/µl together with *unc-122p::tagBFP* (40 ng/µl) to generate INF421 (*syIs390; nonEx108 [nlp-51p::NLS::cGAL(DBD)::cGAL(AD)::let-858 3′UTR + unc-122p::tagBFP]*). We then isolated a spontaneous integrant as INF504 (*syIs390; nonIs11 [nlp-51p::NLS::cGAL(DBD)::cGAL(AD)::let-858 3′UTR + unc-122p::tagBFP]*). For a list of PCR primers used for plasmid constructions, see Table S4.

### Behavioral assays

We performed population chemotaxis assays as described^34^ at 20°C on 9-cm modified chemotaxis agar plates (17 g/L agar, 1 mM CaCl_2_, 1 mM MgSO_4_, 25 mM H_2_KPO_4_/HK_2_PO_4_ pH 6.0) that we poured 3 days before use and stored at room temperature (20-22°C)^32^. To generate a radial H_2_O_2_ gradient, we spotted 20 µL of 300 mM H_2_O_2_ at the “test” position 3 cm from the plate center (Figure S1A) twice—first 22 h and again 3.5 h before the assay—keeping plates at room temperature between applications. We applied an equal volume of autoclaved reverse-osmosis deionized (RODI) water 3 cm from the center at the diametrically opposite “control” position. To generate age-synchronous cohorts of day-1 adults, we placed twenty gravid hermaphrodites on OP50-seeded NGM plates to lay eggs for 3-4 hours, removed the adults, and cultured the eggs for 72-76 hours until the progeny reached the gravid adult stage. We washed these animals three times in M9 buffer and placed them at the center of each chemotaxis plate. For assays with sodium azide, we applied 2 µL of 0.5 M sodium azide to both the test and control spots 10 min before placing the worms. We incubated plates at 20°C for 2 hours and then recorded the number of worms in the test and control regions. We calculated an H_2_O_2_-avoidance index as (number of animals in the control region – number of animals in the test region) / (total number of animals – number of animals at the plate center spot) (Figure S1A). We estimated the H_2_O_2_ concentration along the radial gradient using the diffusion equation for a point source in a 5-mm-deep cylindrical aqueous volume^133^, using an H_2_O_2_ diffusion coefficient of 1.5 x 10^−5^ cm^2^ sec^−1^ (Ref. ^134^).

### Microscopy

Chemotaxis plates were illuminated from below with red LED light (∼650 nm; Melpo BLFL-LFFA) diffused through translucent plastic inside an opaque enclosure. Animals were imaged from above at 120 frames sec^−1^ for 45 minutes using a FLIR Grasshopper 3 camera (1200 × 1200 pixels). To reduce noise and facilitate downstream processing, frames were averaged in 1-second intervals during acquisition. The mean frame over the full recording was then subtracted from each individual frame to enhance motion contrast. To generate track figures and videos, brightness-scaled, average-subtracted frames were projected over a 20-frame window using the Wormtrails python package^135^. Before mean-frame subtraction, vignetting was corrected by dividing each frame by a Gaussian-blurred version of the average frame with a 100-pixel standard deviation.

### Survival assays

We conducted lifespan assays as described^136^, with the following modifications. We cultured nematodes on *E. coli* JI377, which does not degrade H_2_O_2_ in the environment^32,36^. At the onset of adulthood, we transferred worms to plates containing 10 µg/mL 5-fluoro-2′-deoxyuridine (FUDR) to prevent vulval rupture^137^ and eliminate live progeny. For assays with H_2_O_2_, we added the compound to the molten agar immediately before pouring. We used the L4 molt as age 0 for lifespan analysis. We assessed viability and movement daily until death, as described^138^.

### Calcium imaging

Calcium imaging experiments were performed as previously described^21^, with some modifications. We picked young adult animals, placed them in a microfluidic device that delivered stimuli alternating with buffer^21,139^. For H_2_O_2_ dose-response experiments, we used a microfluidic device capable of delivering up to 13 different stimuli to a single immobilized animal^21,139^. We delivered each stimulus for 15 seconds, separated by buffer (H_2_O) for 45 seconds. Each animal was presented twice with a set of stimuli. Fluorescence was recorded with a spinning disc confocal microscope (Dragonfly 200, Andor) and a sCMOS camera (Kinetix, Teledyne Photometrics) that captured fluorescence from GCaMP6s at 15 ms/1.1 µm z-slice, 25 z-slices/volume, and 2.67 volumes/second. To image all 14 pharyngeal neuron classes, as well as ASK and RIP, we used *otIs672[rab-3p::NLS::GCaMP6s + arrd-4::NLS:::GCaMP6s]*^140^. To image ADF, ADL, AFD, ASE, ASG, ASH, ASI, ASJ, ASK, AWA, AWB, AWC, BAG, IL2V, and URX we used *aeaIs8[ift-20p::GCaMP6s::3xNLS + lin-15(+)]*^141^. Neuronal nuclei were identified according to published anatomical maps and manuals^140,141^. To image the AIA neurite, we used *syEx1793[sra-9p::GCaMP6s, sre-1p::GCaMP6s, gcy-28.dp::GCaMP6s]*^142^, which drives GCaMP6s expression in the soma and processes of ASK, ADL, and AIA. When visible, ASK and ADL were identified on the dorsal side, and AIA on the ventral side; the AIA process appeared as a distinct punctum near the soma. Neuron identities were further confirmed using a positive control stimulus (bacterial OP50 suspension), which inhibits ASK and excites ADL and AIA^142^. To extract calcium activity from neuronal nuclei, we identified the center of each neuronal nucleus in every frame and took the average pixel intensities of a 3.2 µm x 3.2 µm x 3.6 µm rectangular box around those centers; the same approach was used for the AIA neurite by identifying a point on the process immediately adjacent to the soma in every frame. The neuron-independent background signal was removed and F/F_0_ calculated for each stimulus-response, where F_0_ was the intensity of that neuron 3.75 seconds before the stimulus was delivered.

## Statistical analysis

We performed all statistical analyses in JMP Pro 18 (SAS). Survival curves were calculated using the Kaplan-Meier method. We used the log-rank test to determine if the survival functions of two or more groups were equal. We compared group means for chemotaxis indices and GCaMP6 fluorescence by ANOVA. When more than two groups were present, we used Tukey’s HSD test to identify pairwise differences. To test interactions, we fit ordinal or least-squares regression models of the form response = intercept + group 1 + group 2 + group 1 * group 2*+ ε*. The second to last term in this model quantifies the existence, magnitude, and type (synergistic or antagonistic) of interaction between groups. We adjusted *P* values for multiple testing with the Bonferroni method.

## Materials availability

Further information and requests for resources and reagents should be directed to and will be fulfilled by the corresponding authors Javier Apfeld and Vivek Venkatachalam.

## Acknowledgements

We thank Nikhil Bhatla for detailed comments on our manuscript. Joy Alcedo, Cori Bargmann, Rita Droste, Miriam Goodman, Joshua Kaplan, Takaaki Hirotsu, H. Robert Horvitz, Ikue Mori, Junho Lee, Roger Pocock, Douglas Portman, and Shawn Xu kindly provided strains. Chen Wang and Oliver Hobert kindly gave permission to modify their diagram of the *C. elegans* hermaphrodite nervous system. Some of the plasmid constructs were made by Dr. Wolfgang Bönigk from the facility of genetic engineering at MPINB. The I1 driver construct was injected by SUNY Biotech, CN. pHW393, pHW530, and pJL081 are gifts from Han Wang and Paul Sternberg. We benefited from discussions with members of Javier Apfeld’s, Erin Cram’s, and Vivek Venkatachalam’s labs, Peter Askjaer, Nikhil Bhatla, Steven Cook, Max Heiman, Oliver Hobert, H. Robert Horvitz, Steven Sando, and Paul Sternberg. We derived some information from Wormbase, which is supported by the National Human Genome Research Institute at the NIH (grant #U41 HG002223), the UK Medical Research Council, and the UK Biotechnology and Biological Sciences Research Council. Some strains were provided by the CGC, which is funded by NIH Office of Research Infrastructure Programs (P40 OD010440). J.A. acknowledges support from a National Science Foundation CAREER grant #1750065, a Hevolution Foundation GRO award, and a Longevity Impetus grant from Norn Group, Hevolution Foundation and Rosenkranz Foundation. V.V. acknowledges support from the NIH (R01 NS126334 and R01 DK142077). Part of this work was funded through the BaBots project to M.S and J.L. The BABots project has received funding from the Horizon Europe, PathFinder European Innovation Council Work Programme under grant agreement No 101098722. Views and opinions expressed are, however, those of the authors only and do not necessarily reflect those of the European Union or European Innovation Council and SMEs Executive Agency (EISMEA). Neither the European Union nor the granting authority can be held responsible for them.

## Competing interests

The authors declare that no competing interests exist.

## Figure Legends

**Figure S1.**
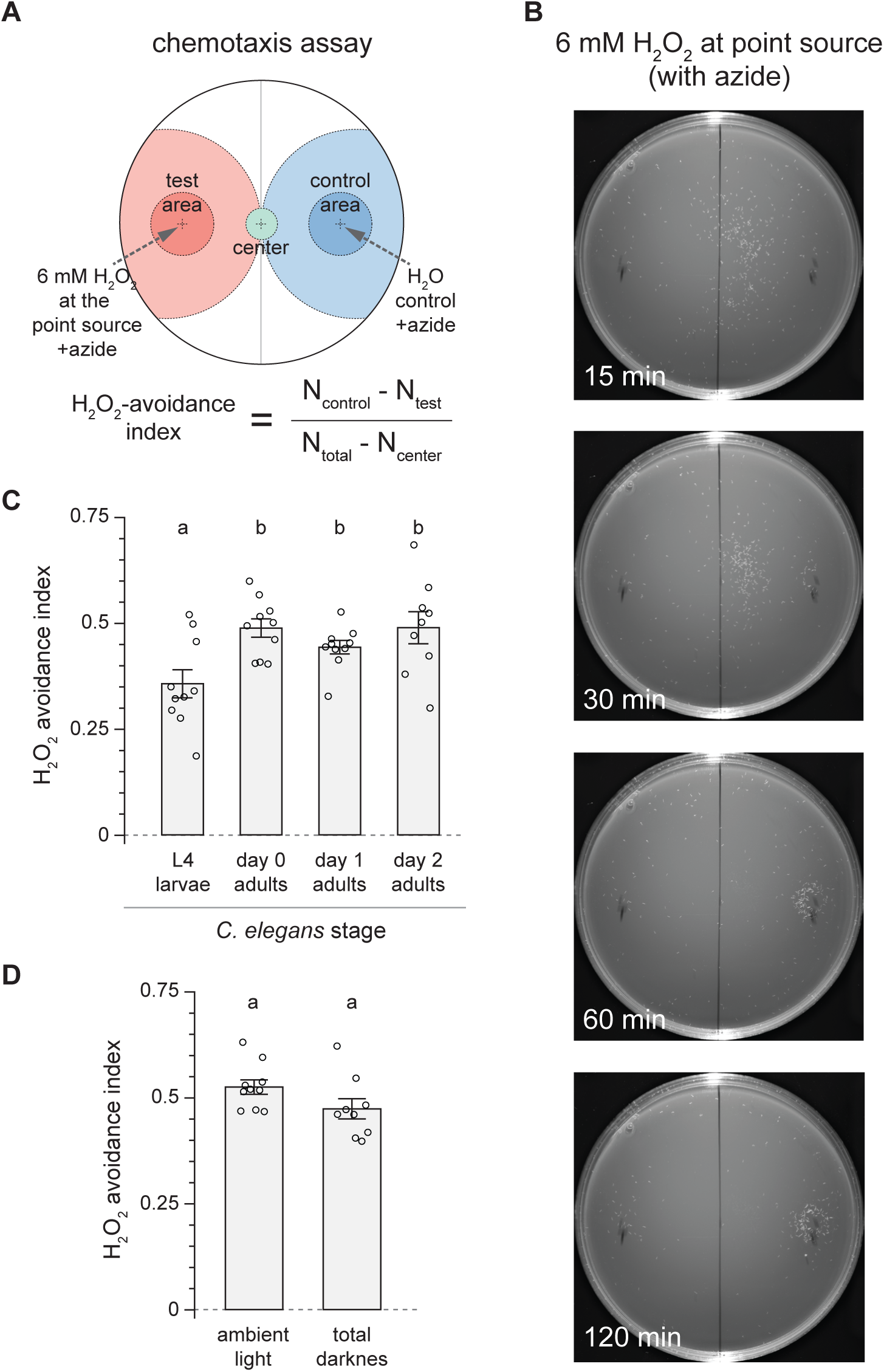
*C. elegans* chemotaxes away from non-lethal concentrations of H_2_O_2_. (A) Schematic of the chemotaxis assay. *C. elegans* were placed at the center (green) of an agar plate containing an H_2_O_2_ gradient emanating from a point source in the test area (red) and allowed to crawl freely. Ten minutes before adding the worms, sodium azide was added at the H_2_O_2_ point source and at a diametrically opposite “control” spot to immobilize animals upon arrival. The H_2_O_2_-avoidance index was calculated as indicated based on the number of animals in each region at the end of the assay. (B) Representative photographs of day-1 wild-type *C. elegans* during an H_2_O_2_-avoidance assay with 6 mM H_2_O_2_ at the point source and the paralytic agent sodium azide at the point-source and control areas, captured at the indicated time points. (C) L4 larvae displayed a modest decrease in H_2_O_2_ avoidance compared with adults. (D) H_2_O_2_ avoidance was not affected by ambient light. Data in (C-D) are plotted as mean ± s.e.m. Groups labeled with different letters exhibited significant differences (*P* < 0.05, Tukey HSD test), otherwise not significant (*P* > 0.05).

**Figure S2.**
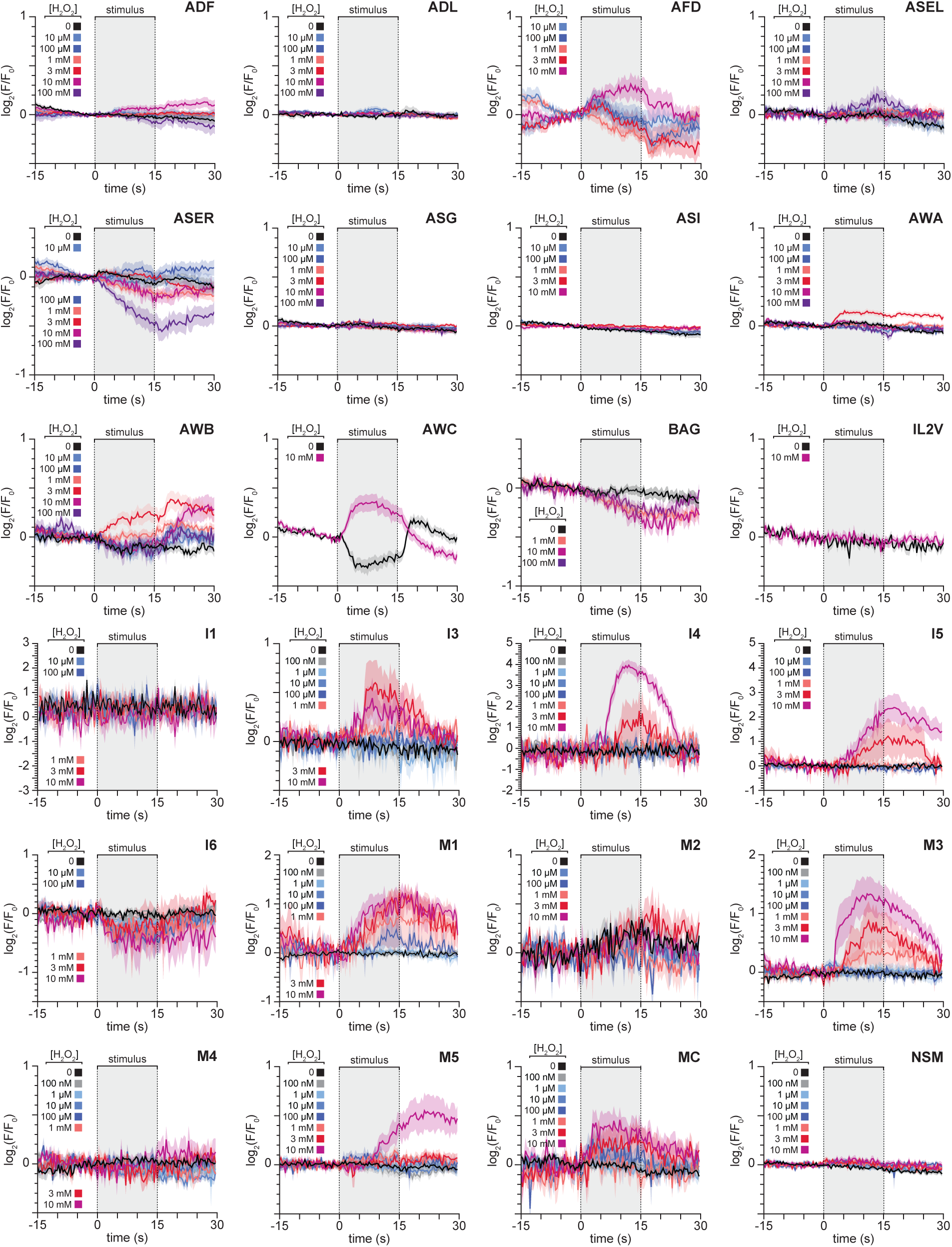
Sensory neurons in both the somatic and pharyngeal nervous systems exhibit neuron-specific, dose-dependent responses to H_2_O_2_. Mean GCaMP6 fluorescence traces for specific neuronal classes in response to the indicated H_2_O_2_ concentrations. Data are plotted as mean ± s.e.m. The shaded box marks the stimulus delivery interval. The remaining neuronal classes are shown in Figure 3. Statistical analyses for all panels are in Table S6.

**Figure S3:**
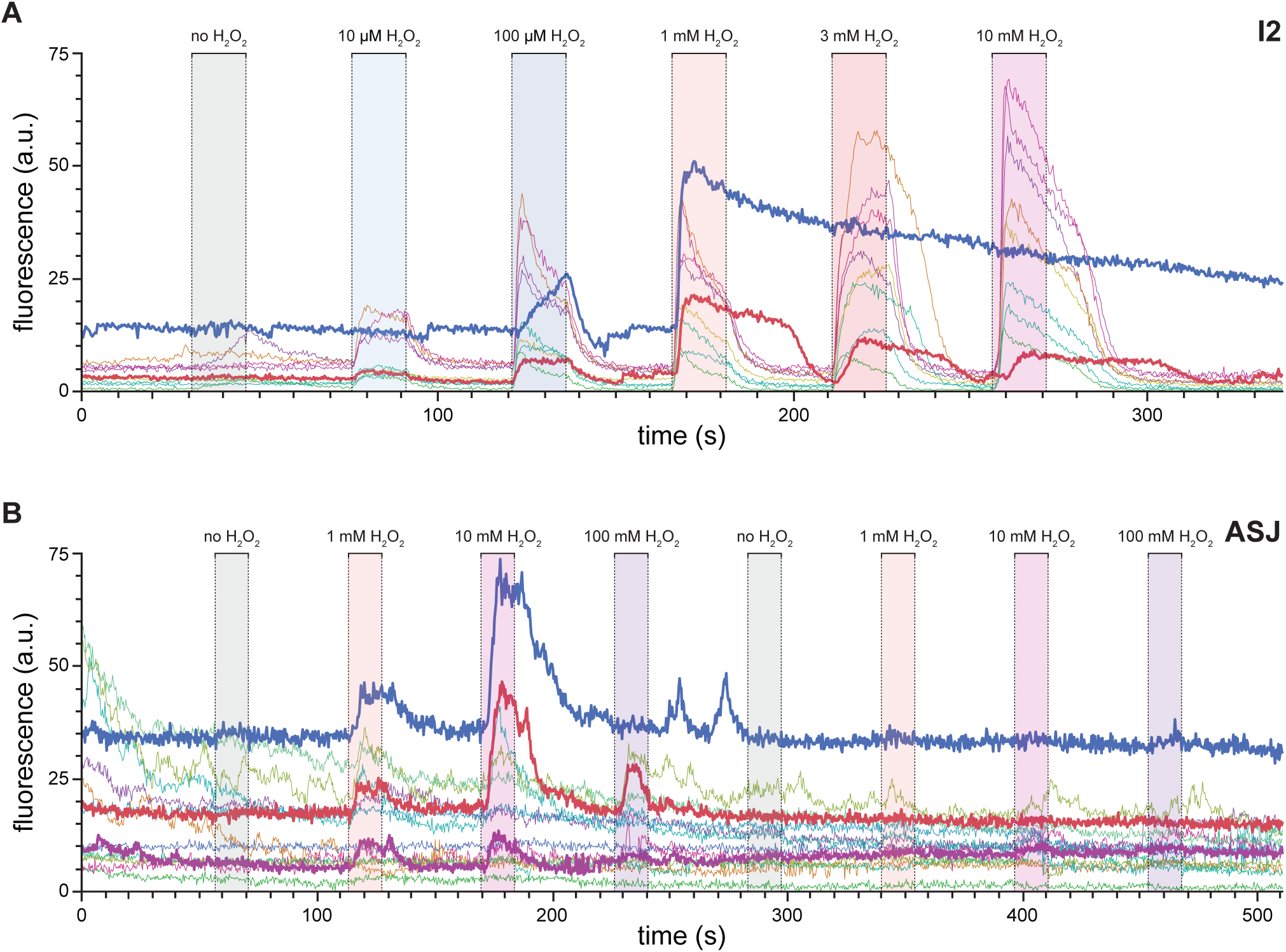
H_2_O_2_ concentration and exposure history can influence neuronal-class response dynamics. (A) Representative GCaMP fluorescence traces from individual I2 neurons during a dose-escalation series with H_2_O_2_. Shaded boxes indicate stimulus delivery at the indicated concentrations. In some neurons (red and blue traces), the 1 mM pulse produced a prolonged OFF response that abolished or strongly attenuated subsequent responses to 3 mM and 10 mM H_2_O_2_. (B) Representative GCaMP fluorescence traces from individual ASJ neurons during two successive dose-escalation series with H_2_O_2_. In some neurons (purple, red, blue), incomplete recovery after high-dose stimulation blunted responses to 100 mM H_2_O_2_ in the first series, and responses in the second series, illustrating history-dependent desensitization.

**Figure S4.**
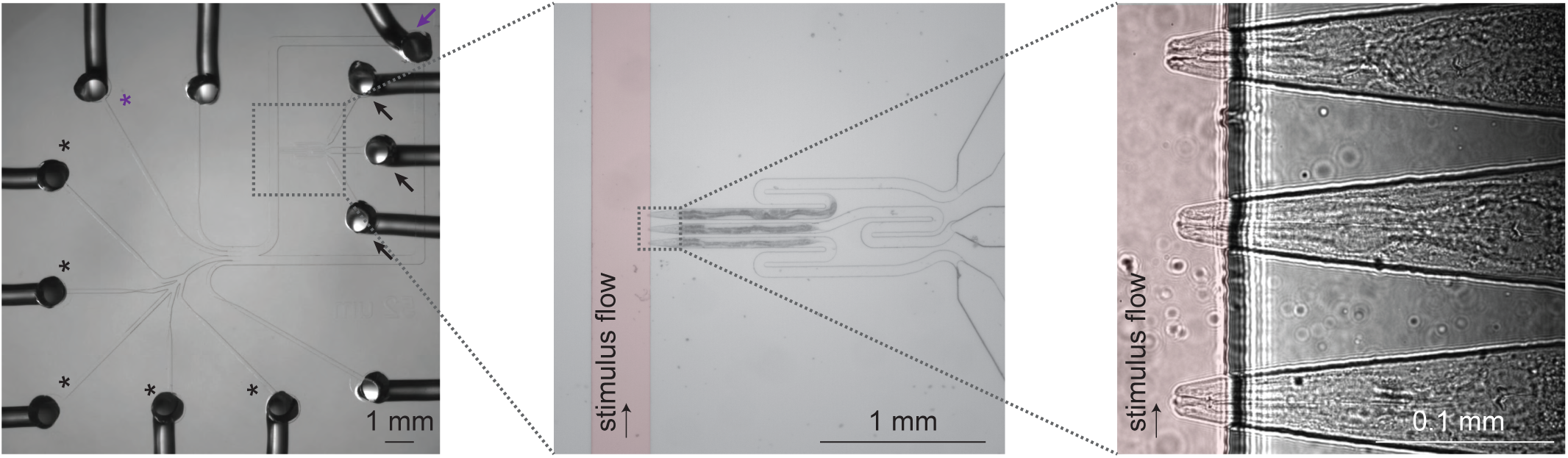
Schematic of the microfluidic platform used for controlled stimulus delivery and simultaneous calcium imaging of three worms. (Left) Image of the multi-channel microfluidic device for multi-worm imaging. Black arrows: worm-loading inlets; purple arrow: waste outlet. Black asterisks: stimulus delivery channels; purple asterisk: buffer channel. (Middle) Enlarged view of the channels where the sensory endings in the nose and mouth of the immobilized nematodes are stimulated with H_2_O_2_. (Right) High-magnification view of the three tapered trapping channels showing individual worms immobilized with their noses and mouths exposed to the stimulus delivery channel; arrow indicates the direction of stimulus flow.

**Figure S5.**
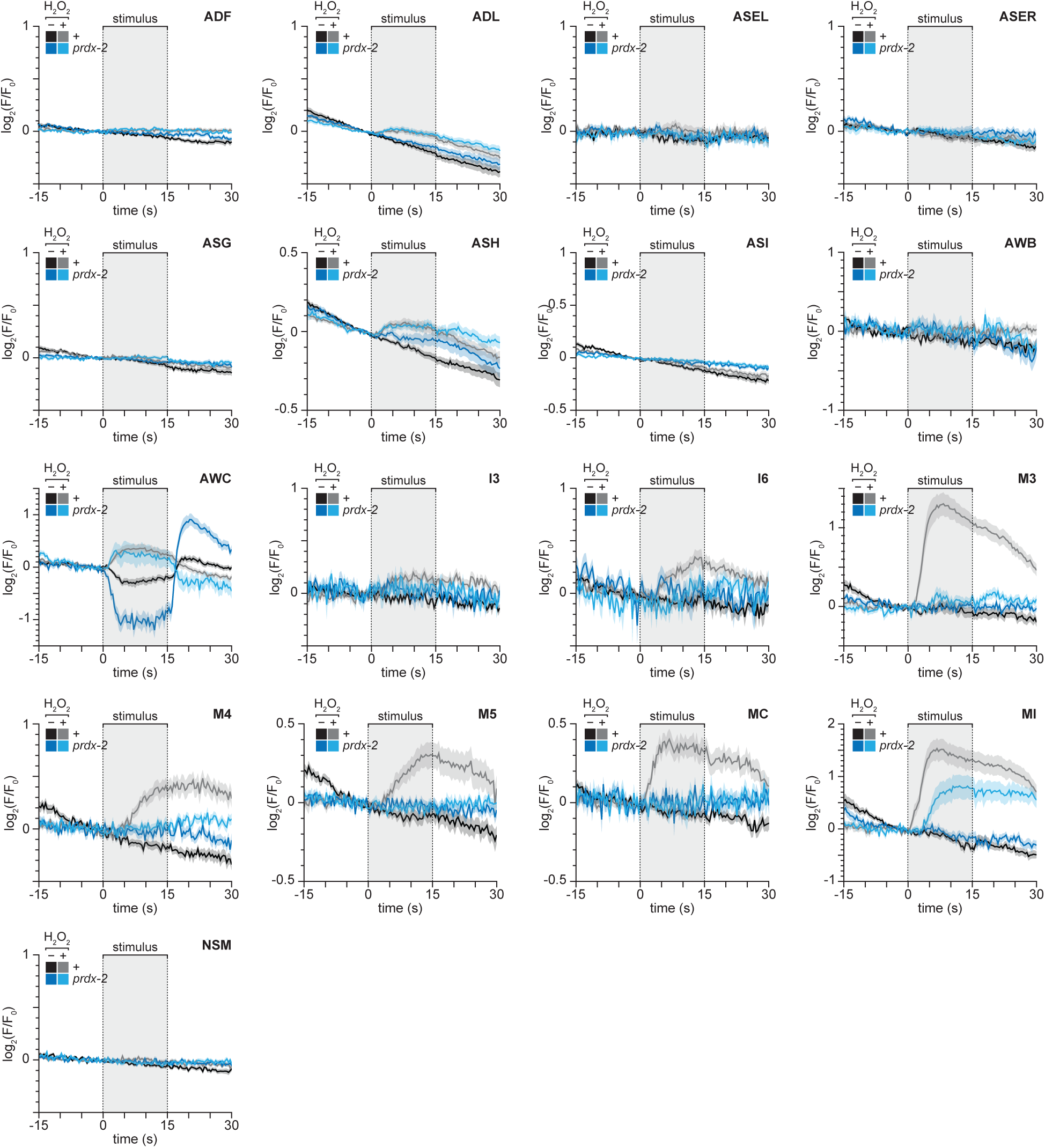
Perception of H_2_O_2_ by sensory neurons in both the somatic and pharyngeal nervous systems requires the cytosolic peroxiredoxin PRDX-2. Mean GCaMP6 fluorescence traces for specific neuronal classes in response to 0 and 10 mM H_2_O_2_ in wild type animals and *prdx-2(gk169)* mutants. Data are plotted as mean ± s.e.m. The shaded box marks the stimulus delivery interval. The remaining neuronal classes are shown in Figure 5. Statistical analysis is in Table S7.

**Figure S6.**
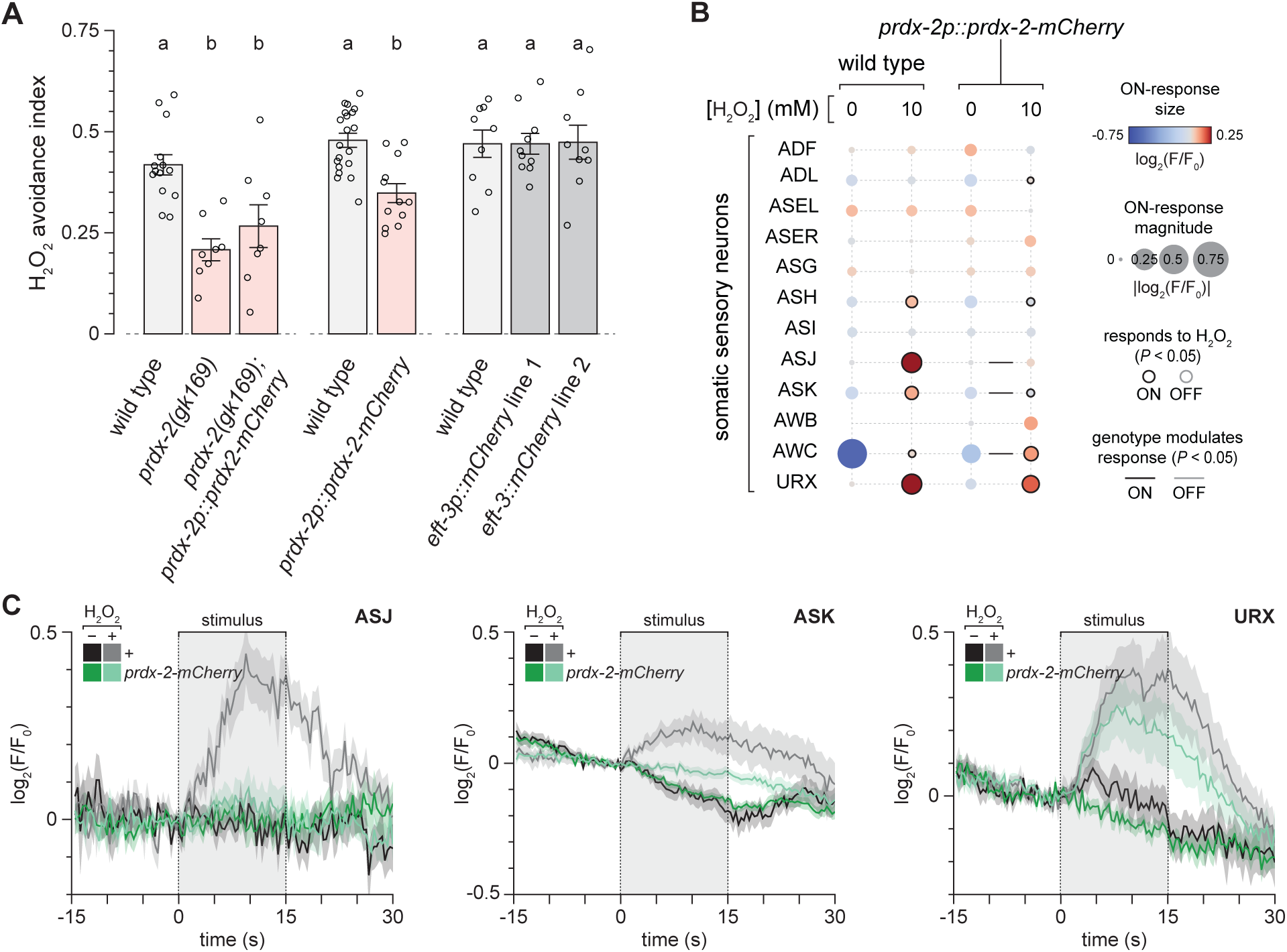
The PRDX-2 C-terminus plays a role in H_2_O_2_ perception and avoidance. (A) A *prdx-2p::prdx-2-mCherry* transgene did not increase H_2_O_2_ avoidance in *prdx-2* mutants but lowered avoidance in an otherwise wild-type background. Two transgenes driving ubiquitous mCherry expression in somatic cells did not affect H_2_O_2_ avoidance. Data are plotted as mean ± s.e.m. Groups labeled with different letters exhibited significant differences (*P* < 0.05, Tukey HSD test), otherwise not significant (*P* > 0.05). (B) Responses of somatic and pharyngeal neuron classes to 0 and 10 mM H_2_O_2_ in wild type animals and in *nEx2078[prdx-2p::prdx-2-mCherry]* transgenics. Each circle represents one neuron class at a given H_2_O_2_ concentration. Circle color encodes the mean change in GCaMP6 fluorescence during the 15 second stimulus interval (ON response); circle area scales with the absolute response magnitude. Black rings indicate ON responses that differ significantly from the no-H_2_O_2_ control (*t*-test, *P* < 0.05); gray rings indicate significant OFF-responses measured during the 15 second post-stimulus interval (*t*-test, *P* < 0.05); unmarked circles indicate responses that were not significant in either interval (*P* > 0.05). Black and grey lines indicate that ON– and OFF-responses, respectively, are affected by the *nEx2078* transgene (standard least-squares regression, *P* < 0.05), otherwise not significant (*P* > 0.05). (C) Mean GCaMP6 fluorescence traces for ASJ, ASK, and URX neuronal classes in response to 0 and 10 mM H_2_O_2_ in wild type animals and in *nEx2078[prdx-2p::prdx-2-mCherry]* transgenics. Data are plotted as mean ± s.e.m. The shaded box marks the stimulus delivery interval. The remaining neuronal classes are shown in Figure S7. Statistical analysis is in Table S9.

**Figure S7.**
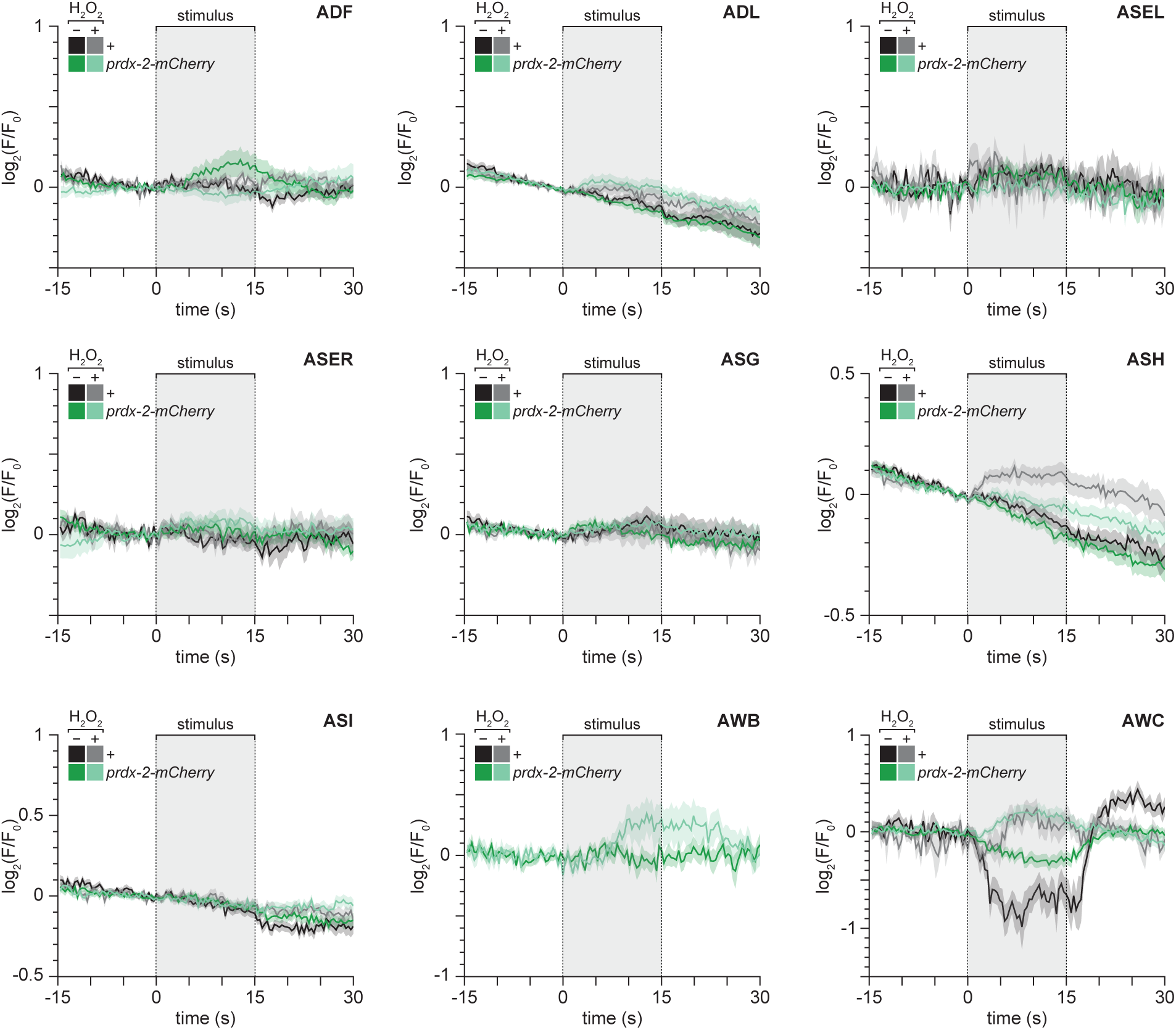
The PRDX-2 C-terminus plays a role in H_2_O_2_ perception and avoidance. Mean GCaMP6 fluorescence traces for specific neuronal classes in response to 0 and 10 mM H_2_O_2_ in wild type animals and in *nEx2078[prdx-2p::prdx-2-mCherry]* transgenics. Data are plotted as mean ± s.e.m. The shaded box marks the stimulus delivery interval. The remaining neuronal classes are shown in Figure S6. Statistical analysis is in Table S9.

**Figure S8.**
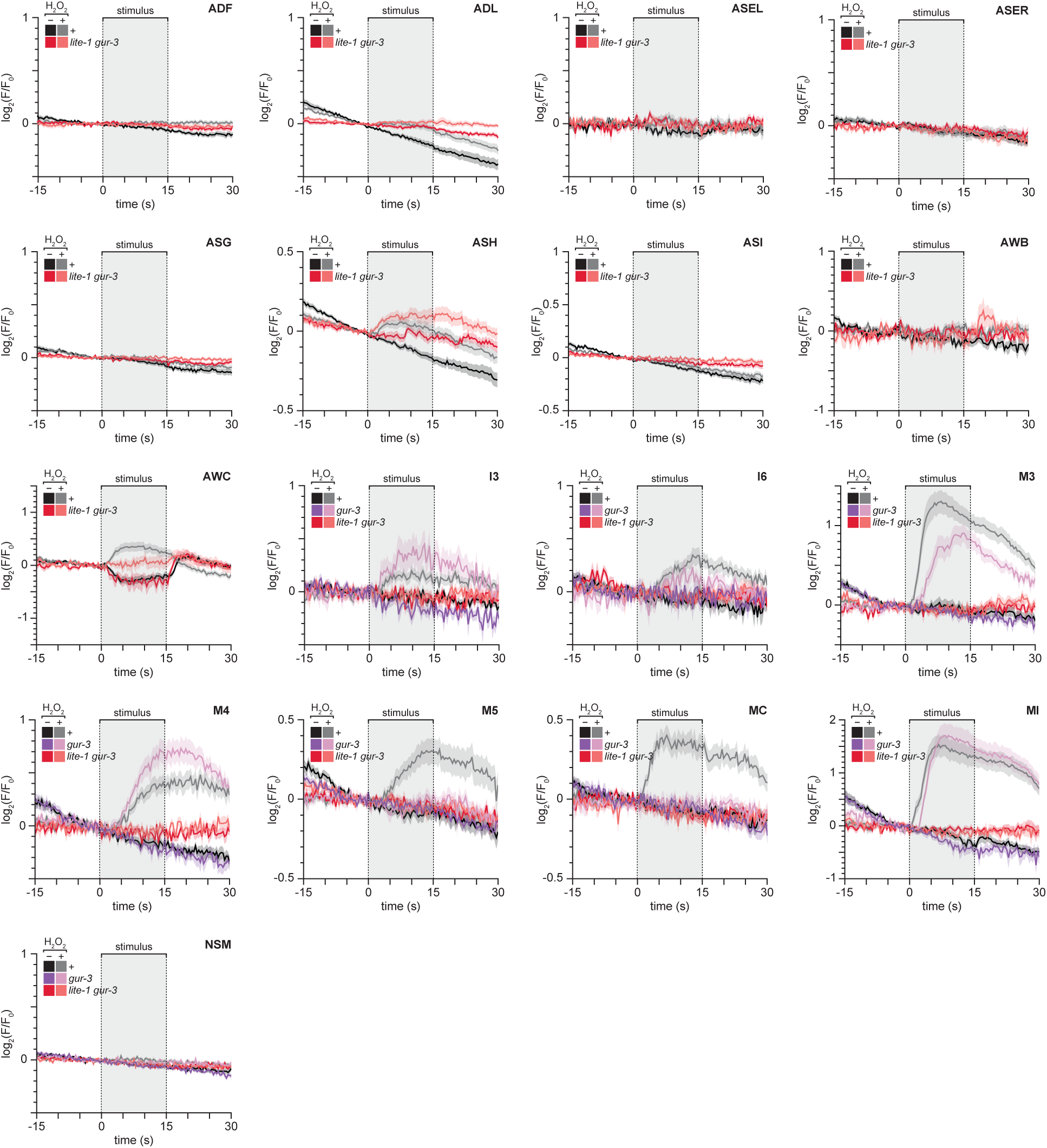
Chemotactic avoidance of H_2_O_2_ is driven by somatic and pharyngeal neuronal sensing through the ion channels LITE-1 and GUR-3. Mean GCaMP6 fluorescence traces for specific neuronal classes in response to 0 and 10 mM H_2_O_2_ in wild-type animals, in *gur-3(ok2245)* mutants, and in *lite-1(ce314) gur-3(ok2245)* double mutants. Data are plotted as mean ± s.e.m. The shaded box marks the stimulus delivery interval. The remaining neuronal classes are shown in Figures 6. Statistical analysis is in Table S10.

**Figure S9.**
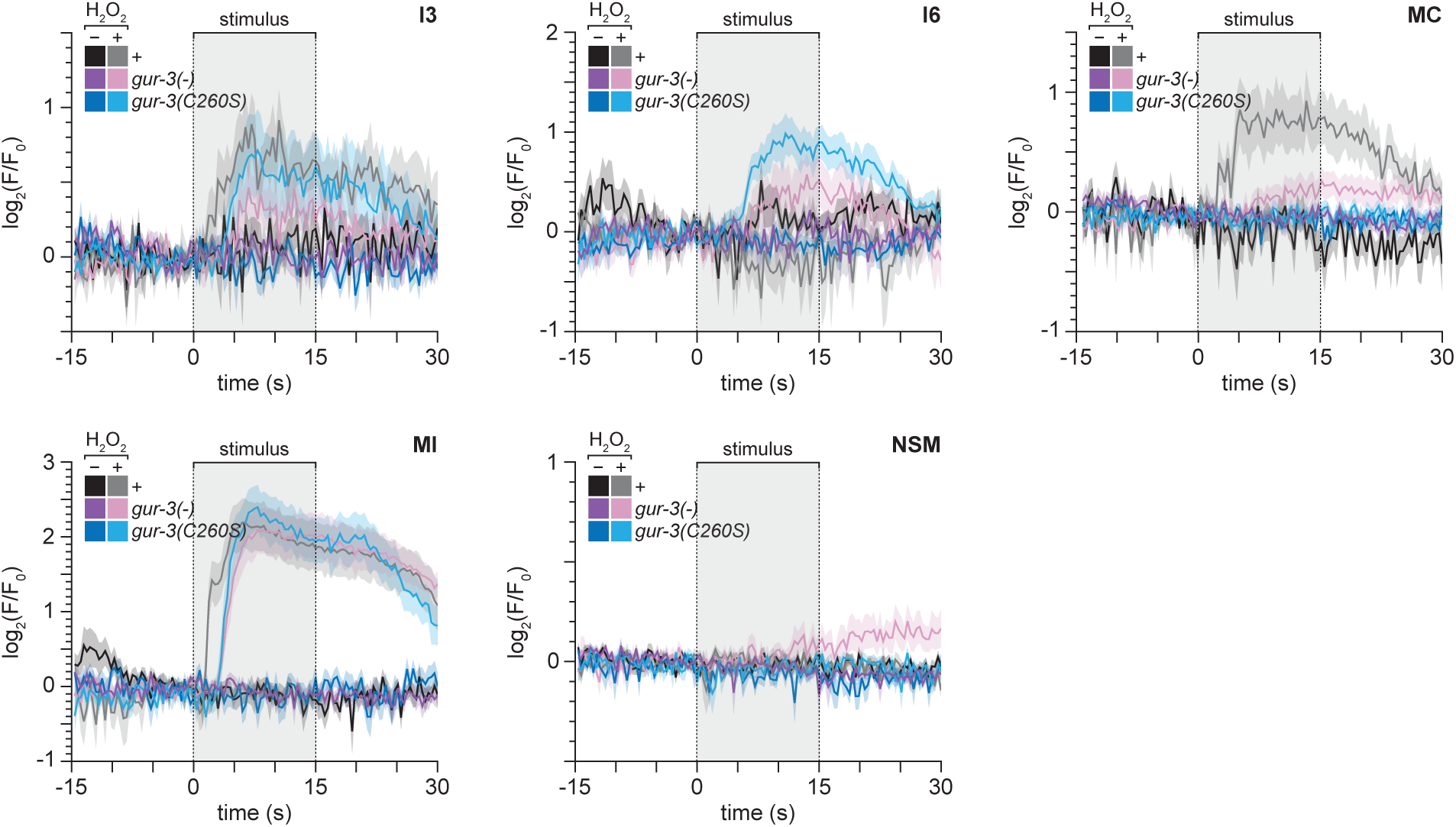
GUR-3 cysteines mediate H_2_O_2_ perception by I2 neurons leading to chemotactic avoidance. Mean GCaMP6 fluorescence traces for specific neuronal classes in response to 0 and 10 mM H_2_O_2_ in wild type animals, in *gur-3(ok2245)* mutants, and in *gur-3(C260S)* mutants. Data are plotted as mean ± s.e.m. The shaded box marks the stimulus delivery interval. The remaining neuronal classes are shown in Figures 7. Statistical analysis is in Table S11.

**Figure S10.**
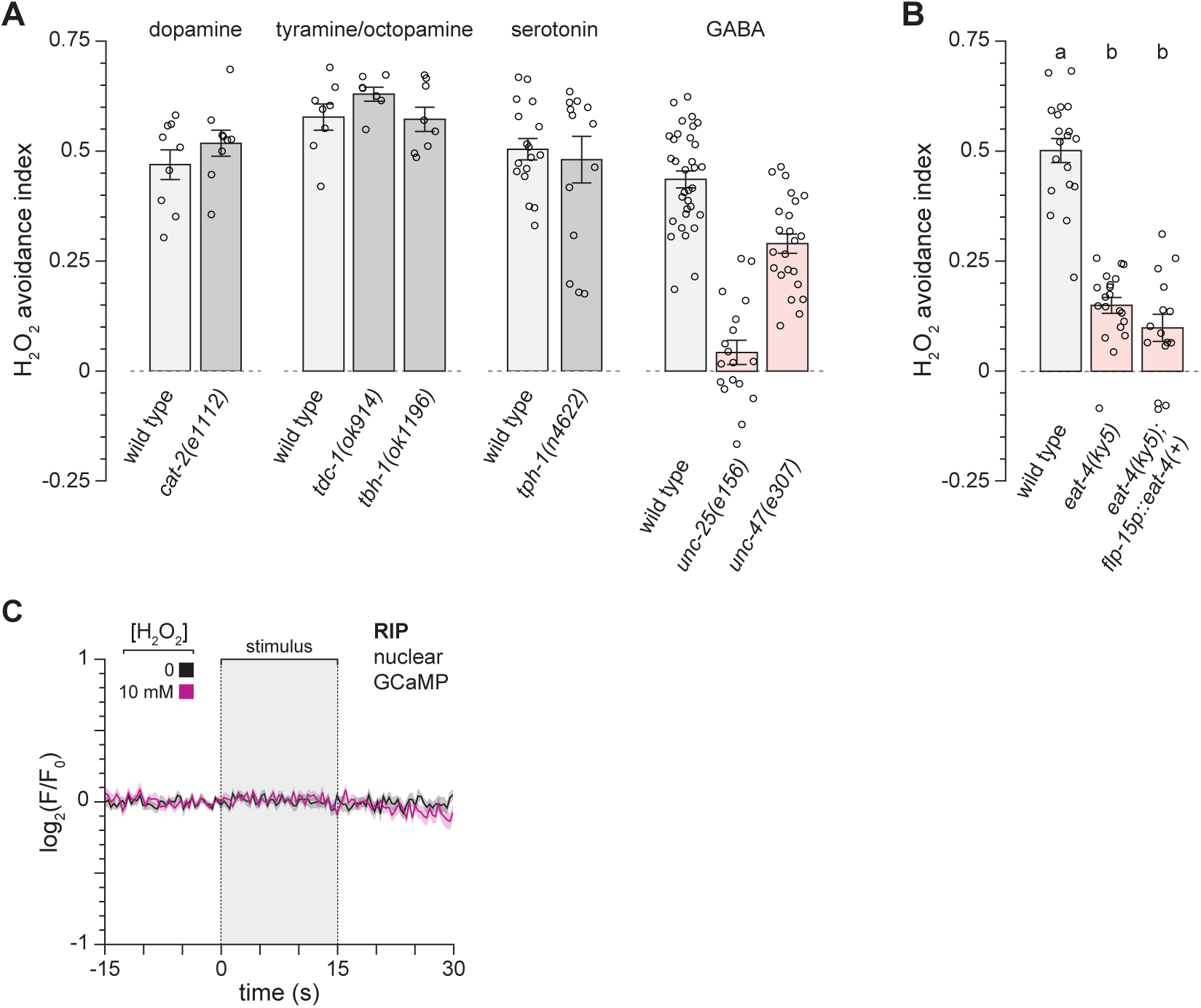
The neurotransmitter GABA promotes avoidance of H_2_O_2_. (A) H_2_O_2_ avoidance in mutants defective in the biosynthesis or transport of specific neurotransmitters. Data are plotted as mean ± s.e.m. Red bars mark groups with a significant decrease relative to the wild-type control in the same panel (ANOVA, *P* < 0.05), otherwise not significant (*P* > 0.05). (B) Expression of *eat-4(+)* in I2 and PHA neurons driven by the *flp-15* promoter did not increase H_2_O_2_ avoidance in *eat-4(ky5)* mutants. Data are plotted as mean ± s.e.m. Groups labeled with different letters exhibited significant differences (*P* < 0.05, Tukey HSD test), otherwise not significant (*P* > 0.05). (C) Mean GCaMP6 fluorescence traces for RIP nuclei in response to the indicated H_2_O_2_ concentrations. The shaded box marks the stimulus delivery interval.

**Supplementary Data 1. Sequences of the wild type and edited alleles**.

## Notes

### Competing Interest Statement

The authors have declared no competing interest.

