## Supplementary data 1 for "The *C. elegans* nervous system reads the internal state of the hydrogen peroxide-detoxification machinery to trigger escape from this common reactive chemical"

>*prdx-2(+)*

ATGTCGAAAGCATTCATCGGAAAGCCAGCTCCACAATTCAAGACTCAAGCCGTCGTTGATGGCGAGTTCGTTGATGTTTCGCTCTCTGACTACAAGGGAAAATACGTTGTGCTCTTCTTCTACCCACTTGACTTCACTTTCGTG**TGC**CCA**ACC**GAGATTATCGCCTTCTCTGACCGTGCTGAGGAGTTCAAGGCTATCAACACCGTTGTGCTCGCCGCTTCCACCGACTCTGTCTTCTCTCACTTGGCATGGATCAACCAGCCACGCAAGCACGGAGGACTCGGAGAGATGAACATTCCAGTTCTCGCTGACACCAACCACCAAATCTCCCGTGATTACGGAGTTCTCAAGGAGGACGAAGGAATTGCTTTCCGTGgtgagttatattctgaatggaatccttcatattttcgtactttatttgaaaaataaaatttcagGACTCTTCATCATCGACCCATCACAAAACCTCCGTCAAATCACCATCAATGATCTTCCAGTCGGACGCTCTGTTGATGAGACTCTTCGTCTTGTTCAGGCCTTCCAGTTCGTCGAGAAGCACGGA**GAG**GTT**TGCCCAGCT**GGATGGACTCCAGGATCCGACACCATCAAGCCAGGAGTCAAAGAAAGCCAAGAGTACTTCAAGAAGCACTAAatgtcttacatctctaatttcccccgtatccctagtatttatctgaacggtttatttatactcatttttcattttgatactgaaattctgaaattcatgaaattgcttgataaccgtaaaataaactcggtattcgaa

Codons for Cysteines 55 and 176 are labeled in **purple**.

A Exon

A Exon

a Intron

a UTR

>*prdx-2(syb10421 C55S)*

ATGTCGAAAGCATTCATCGGAAAGCCAGCTCCACAATTCAAGACTCAAGCCGTCGTTGATGGCGAGTTCGTTGATGTTTCGCTCTCTGACTACAAGGGAAAATACGTTGTGCTCTTCTTCTACCCACTTGACTTCACTTTCGTG**TCA**CCA**ACG**GAGATTATCGCCTTCTCTGACCGTGCTGAGGAGTTCAAGGCTATCAACACCGTTGTGCTCGCCGCTTCCACCGACTCTGTCTTCTCTCACTTGGCATGGATCAACCAGCCACGCAAGCACGGAGGACTCGGAGAGATGAACATTCCAGTTCTCGCTGACACCAACCACCAAATCTCCCGTGATTACGGAGTTCTCAAGGAGGACGAAGGAATTGCTTTCCGTGgtgagttatattctgaatggaatccttcatattttcgtactttatttgaaaaataaaatttcagGACTCTTCATCATCGACCCATCACAAAACCTCCGTCAAATCACCATCAATGATCTTCCAGTCGGACGCTCTGTTGATGAGACTCTTCGTCTTGTTCAGGCCTTCCAGTTCGTCGAGAAGCACGGAGAGGTTTGCCCAGCTGGATGGACTCCAGGATCCGACACCATCAAGCCAGGAGTCAAAGAAAGCCAAGAGTACTTCAAGAAGCACTAAatgtcttacatctctaatttcccccgtatccctagtatttatctgaacggtttatttatactcatttttcattttgatactgaaattctgaaattcatgaaattgcttgataaccgtaaaataaactcggtattcgaa

Missense mutations are labeled in **red**. Synonymous mutations are labeled in **blue**.

A Exon

A Exon

a Intron

a UTR

>*prdx-2(syb10493 C176S)*

ATGTCGAAAGCATTCATCGGAAAGCCAGCTCCACAATTCAAGACTCAAGCCGTCGTTGATGGCGAGTTCGTTGATGTTTCGCTCTCTGACTACAAGGGAAAATACGTTGTGCTCTTCTTCTACCCACTTGACTTCACTTTCGTGTGCCCAAccGAGATTATCGCCTTCTCTGACCGTGCTGAGGAGTTCAAGGCTATCAACACCGTTGTGCTCGCCGCTTCCACCGACTCTGTCTTCTCTCACTTGGCATGGATCAACCAGCCACGCAAGCACGGAGGACTCGGAGAGATGAACATTCCAGTTCTCGCTGACACCAACCACCAAATCTCCCGTGATTACGGAGTTCTCAAGGAGGACGAAGGAATTGCTTTCCGTGgtgagttatattctgaatggaatccttcatattttcgtactttatttgaaaaataaaatttcagGACTCTTCATCATCGACCCATCACAAAACCTCCGTCAAATCACCATCAATGATCTTCCAGTCGGACGCTCTGTTGATGAGACTCTTCGTCTTGTTCAGGCCTTCCAGTTCGTCGAGAAGCACGGA**GAA**GTT**TCACCTGCA**GGATGGACTCCAGGATCCGACACCATCAAGCCAGGAGTCAAAGAAAGCCAAGAGTACTTCAAGAAGCACTAAatgtcttacatctctaatttcccccgtatccctagtatttatctgaacggtttatttatactcatttttcattttgatactgaaattctgaaattcatgaaattgcttgataaccgtaaaataaactcggtattcgaa

Missense mutations are labeled in **red**. Synonymous mutations are labeled in **blue**.

A Exon

A Exon

a Intron

a UTR

>*gur-3(+)*

agtcagtcacgttttcctgctatccaaccacgctttaaactcttccaaaatagattctctaaaaccatctctgaaacttttacttaaaATGACGATAACAGCGTCGAATACGTTGGAGTTCAAATGGACGTCACCAAGGAGCTCGAGATCGTCATTCAgtaagttcactggaatgaacctgtgcagccttttatttttcaacattgtttatagGAACAACGACAGATGCGGAACAGAAAATATCAATTGACATGAGTAATACTTACTGTGATCAGgtaaatgatacagtgatggttcgactctaaaaattactgagatttcgaataactaaaaacagttttttggaatgattaaataagtaaaaattctaaatgtatcaacaatacgtgttgttttgaataattgaacataaaagaaaataagaggagacaaagtgtgttaagttgtgaaattttgactgaagcgacacattttcatcaaagggtaataatatgaaaaccataatagccctacatttttgtgggtgcaggtatatttgagactgcacgcacggtgtgcctagatttatttttagtttcaaatatttttcatatttccgaacgaatagtggaagcagcaaaatgtatttgtagcatcaagtatttctagccattttgcaatttttatattgttcagttctaatgaagattgcagaaaaaaacggtaacaaacgcttattcaagaacaaattgtatatgttcagGTTTTGGGGCCTCTGTACTCCTATATGATGGTGTTGGGACTTAATCACACTCACAGTAGCGCTCGGAATACAATGTTTAAGTGGCCATTGACAATCTACAATTATCTCACTCTAGCCATATTGgtaagctacattatgtcctgatcttcgattaaacaagattgtactttctctgcgtgtgtgtaatacgagatgtttataaaagtttagcccagcaagtggaacattttcaaatttttccagACAGCGGCCACAATTCGACGCATTTCACAAATCAAACAAAAATCGGCAACAAATGAGGAAAAAGATGCTGCTTTTCATGTGCTAAACCCGACGTTTGTGCTCACGTTGTGTCATGCTTTGCTAATGTTCAGgttgtgttttgcgcagcgttaagttggttattaaactcaaacatgaacaaacaaatataacgatacatgaaaatcgcacaaagaatacatggtttttgattcgcgggcataagattgtctagtttgctcaaaagtatttgtatttcgggagataattatatttaaaaaattttgaaatgttatggaagtggtcgtttgatatcaagatcgaacaattttttaaagatatgtgtttcctattagtatggtgtacccataagaattgcaatctctaaacaatatttttaatgatatgaacatttaatgatattcttcattcatttttgaggattccttactaagtcacaaaaaaatcgtcgaattaggtattttctagatgccaattcgaggtttcaaaaagtgtcacacacaagttgaatataccttaataaaaacaagttacttccaatttggaacaaaccatcaaaacttagtattgtagtctcttacagtctaatagtattccacgaaaacatttccgggaattcgtatggcacgcaatactattagaaggtaaatcaattttcataaattataaaccaaaaaaaaatattgatggaaaattattgtttgaacgtttctgtagtctattttattaatttattgttttatctcgtatatgacttgtttaattgacttgcattactagacagtctcaatttcctaaaaacgaactagtttctaattttttgaataagactttagtttcatggaaattaaggtttattggaaaccaatcacaaagctaacatcacaaatttattttaaatgatatgcgctttgtgttattctcatatttaaatttcagTGGTCTTGCTGCCGGATTCCTTTTGCTAAAGTTGCAAAAGCAAAGAGAGAAAATGTACCACgtgagtgaaattagacacggttgaggtagatcacaagaccaacagggcttgacttacagGTGTTAGATCAAGGGTTGGGTAGGAATAGAAATGAAGAACATGACAGTCATCATTTCAAGCTGAATAAGTTATTTATATCAATCAGTTTTTCGTTTGCAGCTGCATTATCCTTTGTACAAATTGgtaacaaaaaagtatgactaattcgaaaccattcgaaaaatttccagCTACAAAAATGAGATATCTTGATCTCCCTGACACTCCTGATCTGATTAACAGAAAAATCTACTTTGTGATTCTTGAAGGATACGTGgtacggttgatattctttatgcttgccaaaaattttcaaaattaagATTTTCATCGCTTCCAGTTGCATTTCATTAGTCGCAATATTATTTTTTCAACTATGTAGAATACTCCAATTTTCGATAGGACAACTAATAGAAGAAATGGTGCCTAAAGAAAAAGAAGAA**TGC**CCACTA**CCAGAA**CAATCTTTACAACAAATTCACGATGTTCAAATTCATTACCAAGAAATTTCCAATGCTAAGCTGTATATAGAACAGAATTTCTCATTTTCACTGTTTTATACCTACGGC**TGTTGT**ATTCCG**CTG**ACCTGTCTGTTGGGCTACATTGCTTTTCGAAATGGAATTCAGgtagtgaaaatgagcagcaataccattgtttcatatcatcatttagGCTGATATGGCGGAAACTTTTTCTGTGGCAATATGGCTCACTAATACTATGTTAGCATTGATGCTTTTCTCGATTCCAGCTTTTATGATTGCTGAAGAGgttagtacttataagtatgtatatgtttgtttaaatacatagatatagttagagcagaactgagaaaattgagttagtaaagtttttgtatttcttcaacttttgatgttagctgaacaaaaattaaaaaacaacataaattaaatataatgcaagcatatttgtgaacaagaaatgtgtacgtggataaagtcgcggtggaattatcatttaaccgtgattgcatgcatataaatcccacaatacgtaaactcaatttatttatgacagtttcacaattacgatcagaaatttaaaataaataggttttcaaacactattcttcagcgacaatatttctgctaattgtcgctgatattttatgttcgcaaacgcagtaattcaaaattgatatccaaacattaagttgtacaaataaaaattaataattacagGGCGACAAACTTCTAACCGCTTCTTTCAAAATGTATCACGAAACTCTATGTGAGGAACGTGATCTTCTAGTTTTGgtgtgttcgttatctcaaaaaaaaatcgatcaattctactttcagTCTCAAATGAGTTTTCTTTCGTTTCAAATGCATGCTACAAAGCTTACTCTCACCGCCGGGAATTTCTTCATGATGAATCGCAAAATTATGATCAGTgtacgtttaaaggcgtgcagctggtttttagttgatatttaaaatatttcagTTGTTCTCTGCAATTTTCACATATTTTCTTATTTTGGTACAGTTTGATGCTGAAAAGGAGCGAGCCGGAGAATGCAATAATCAATCTCGTGTGCTCATTGTCCAACCACCTGTGTAAataaaattgtcttaacattttccccatattgagccaacttcaattagtttccaatatgttcatttgtaactacgcaatgtatttgacaaattgtttattcatgtctcctaaatacaagaaagaaagtcattt

Codons for Cysteines 260, 301, and 302 are labeled in **purple**.

A Exon

A Exon

a Intron

a UTR

>*gur-3(syb8201 C260S)*

agtcagtcacgttttcctgctatccaaccacgctttaaactcttccaaaatagattctctaaaaccatctctgaaacttttacttaaaATGACGATAACAGCGTCGAATACGTTGGAGTTCAAATGGACGTCACCAAGGAGCTCGAGATCGTCATTCAgtaagttcactggaatgaacctgtgcagccttttatttttcaacattgtttatagGAACAACGACAGATGCGGAACAGAAAATATCAATTGACATGAGTAATACTTACTGTGATCAGgtaaatgatacagtgatggttcgactctaaaaattactgagatttcgaataactaaaaacagttttttggaatgattaaataagtaaaaattctaaatgtatcaacaatacgtgttgttttgaataattgaacataaaagaaaataagaggagacaaagtgtgttaagttgtgaaattttgactgaagcgacacattttcatcaaagggtaataatatgaaaaccataatagccctacatttttgtgggtgcaggtatatttgagactgcacgcacggtgtgcctagatttatttttagtttcaaatatttttcatatttccgaacgaatagtggaagcagcaaaatgtatttgtagcatcaagtatttctagccattttgcaatttttatattgttcagttctaatgaagattgcagaaaaaaacggtaacaaacgcttattcaagaacaaattgtatatgttcagGTTTTGGGGCCTCTGTACTCCTATATGATGGTGTTGGGACTTAATCACACTCACAGTAGCGCTCGGAATACAATGTTTAAGTGGCCATTGACAATCTACAATTATCTCACTCTAGCCATATTGgtaagctacattatgtcctgatcttcgattaaacaagattgtactttctctgcgtgtgtgtaatacgagatgtttataaaagtttagcccagcaagtggaacattttcaaatttttccagACAGCGGCCACAATTCGACGCATTTCACAAATCAAACAAAAATCGGCAACAAATGAGGAAAAAGATGCTGCTTTTCATGTGCTAAACCCGACGTTTGTGCTCACGTTGTGTCATGCTTTGCTAATGTTCAGgttgtgttttgcgcagcgttaagttggttattaaactcaaacatgaacaaacaaatataacgatacatgaaaatcgcacaaagaatacatggtttttgattcgcgggcataagattgtctagtttgctcaaaagtatttgtatttcgggagataattatatttaaaaaattttgaaatgttatggaagtggtcgtttgatatcaagatcgaacaattttttaaagatatgtgtttcctattagtatggtgtacccataagaattgcaatctctaaacaatatttttaatgatatgaacatttaatgatattcttcattcatttttgaggattccttactaagtcacaaaaaaatcgtcgaattaggtattttctagatgccaattcgaggtttcaaaaagtgtcacacacaagttgaatataccttaataaaaacaagttacttccaatttggaacaaaccatcaaaacttagtattgtagtctcttacagtctaatagtattccacgaaaacatttccgggaattcgtatggcacgcaatactattagaaggtaaatcaattttcataaattataaaccaaaaaaaaatattgatggaaaattattgtttgaacgtttctgtagtctattttattaatttattgttttatctcgtatatgacttgtttaattgacttgcattactagacagtctcaatttcctaaaaacgaactagtttctaattttttgaataagactttagtttcatggaaattaaggtttattggaaaccaatcacaaagctaacatcacaaatttattttaaatgatatgcgctttgtgttattctcatatttaaatttcagTGGTCTTGCTGCCGGATTCCTTTTGCTAAAGTTGCAAAAGCAAAGAGAGAAAATGTACCACgtgagtgaaattagacacggttgaggtagatcacaagaccaacagggcttgacttacagGTGTTAGATCAAGGGTTGGGTAGGAATAGAAATGAAGAACATGACAGTCATCATTTCAAGCTGAATAAGTTATTTATATCAATCAGTTTTTCGTTTGCAGCTGCATTATCCTTTGTACAAATTGgtaacaaaaaagtatgactaattcgaaaccattcgaaaaatttccagCTACAAAAATGAGATATCTTGATCTCCCTGACACTCCTGATCTGATTAACAGAAAAATCTACTTTGTGATTCTTGAAGGATACGTGgtacggttgatattctttatgcttgccaaaaattttcaaaattaagATTTTCATCGCTTCCAGTTGCATTTCATTAGTCGCAATATTATTTTTTCAACTATGTAGAATACTCCAATTTTCGATAGGACAACTAATAGAAGAAATGGTGCCTAAAGAAAAAGAAGAA**TCA**CCACTA**CCGGAG**CAATCTTTACAACAAATTCACGATGTTCAAATTCATTACCAAGAAATTTCCAATGCTAAGCTGTATATAGAACAGAATTTCTCATTTTCACTGTTTTATACCTACGGCTGTTGTATTCCGCTGACCTGTCTGTTGGGCTACATTGCTTTTCGAAATGGAATTCAGgtagtgaaaatgagcagcaataccattgtttcatatcatcatttagGCTGATATGGCGGAAACTTTTTCTGTGGCAATATGGCTCACTAATACTATGTTAGCATTGATGCTTTTCTCGATTCCAGCTTTTATGATTGCTGAAGAGgttagtacttataagtatgtatatgtttgtttaaatacatagatatagttagagcagaactgagaaaattgagttagtaaagtttttgtatttcttcaacttttgatgttagctgaacaaaaattaaaaaacaacataaattaaatataatgcaagcatatttgtgaacaagaaatgtgtacgtggataaagtcgcggtggaattatcatttaaccgtgattgcatgcatataaatcccacaatacgtaaactcaatttatttatgacagtttcacaattacgatcagaaatttaaaataaataggttttcaaacactattcttcagcgacaatatttctgctaattgtcgctgatattttatgttcgcaaacgcagtaattcaaaattgatatccaaacattaagttgtacaaataaaaattaataattacagGGCGACAAACTTCTAACCGCTTCTTTCAAAATGTATCACGAAACTCTATGTGAGGAACGTGATCTTCTAGTTTTGgtgtgttcgttatctcaaaaaaaaatcgatcaattctactttcagTCTCAAATGAGTTTTCTTTCGTTTCAAATGCATGCTACAAAGCTTACTCTCACCGCCGGGAATTTCTTCATGATGAATCGCAAAATTATGATCAGTgtacgtttaaaggcgtgcagctggtttttagttgatatttaaaatatttcagTTGTTCTCTGCAATTTTCACATATTTTCTTATTTTGGTACAGTTTGATGCTGAAAAGGAGCGAGCCGGAGAATGCAATAATCAATCTCGTGTGCTCATTGTCCAACCACCTGTGTAAataaaattgtcttaacattttccccatattgagccaacttcaattagtttccaatatgttcatttgtaactacgcaatgtatttgacaaattgtttattcatgtctcctaaatacaagaaagaaagtcattt

Missense mutations are labeled in **red**. Synonymous mutations are labeled in **blue**.

A Exon

A Exon

a Intron

a UTR

>*gur-3(syb8200 C301S C302S)*

agtcagtcacgttttcctgctatccaaccacgctttaaactcttccaaaatagattctctaaaaccatctctgaaacttttacttaaaATGACGATAACAGCGTCGAATACGTTGGAGTTCAAATGGACGTCACCAAGGAGCTCGAGATCGTCATTCAgtaagttcactggaatgaacctgtgcagccttttatttttcaacattgtttatagGAACAACGACAGATGCGGAACAGAAAATATCAATTGACATGAGTAATACTTACTGTGATCAGgtaaatgatacagtgatggttcgactctaaaaattactgagatttcgaataactaaaaacagttttttggaatgattaaataagtaaaaattctaaatgtatcaacaatacgtgttgttttgaataattgaacataaaagaaaataagaggagacaaagtgtgttaagttgtgaaattttgactgaagcgacacattttcatcaaagggtaataatatgaaaaccataatagccctacatttttgtgggtgcaggtatatttgagactgcacgcacggtgtgcctagatttatttttagtttcaaatatttttcatatttccgaacgaatagtggaagcagcaaaatgtatttgtagcatcaagtatttctagccattttgcaatttttatattgttcagttctaatgaagattgcagaaaaaaacggtaacaaacgcttattcaagaacaaattgtatatgttcagGTTTTGGGGCCTCTGTACTCCTATATGATGGTGTTGGGACTTAATCACACTCACAGTAGCGCTCGGAATACAATGTTTAAGTGGCCATTGACAATCTACAATTATCTCACTCTAGCCATATTGgtaagctacattatgtcctgatcttcgattaaacaagattgtactttctctgcgtgtgtgtaatacgagatgtttataaaagtttagcccagcaagtggaacattttcaaatttttccagACAGCGGCCACAATTCGACGCATTTCACAAATCAAACAAAAATCGGCAACAAATGAGGAAAAAGATGCTGCTTTTCATGTGCTAAACCCGACGTTTGTGCTCACGTTGTGTCATGCTTTGCTAATGTTCAGgttgtgttttgcgcagcgttaagttggttattaaactcaaacatgaacaaacaaatataacgatacatgaaaatcgcacaaagaatacatggtttttgattcgcgggcataagattgtctagtttgctcaaaagtatttgtatttcgggagataattatatttaaaaaattttgaaatgttatggaagtggtcgtttgatatcaagatcgaacaattttttaaagatatgtgtttcctattagtatggtgtacccataagaattgcaatctctaaacaatatttttaatgatatgaacatttaatgatattcttcattcatttttgaggattccttactaagtcacaaaaaaatcgtcgaattaggtattttctagatgccaattcgaggtttcaaaaagtgtcacacacaagttgaatataccttaataaaaacaagttacttccaatttggaacaaaccatcaaaacttagtattgtagtctcttacagtctaatagtattccacgaaaacatttccgggaattcgtatggcacgcaatactattagaaggtaaatcaattttcataaattataaaccaaaaaaaaatattgatggaaaattattgtttgaacgtttctgtagtctattttattaatttattgttttatctcgtatatgacttgtttaattgacttgcattactagacagtctcaatttcctaaaaacgaactagtttctaattttttgaataagactttagtttcatggaaattaaggtttattggaaaccaatcacaaagctaacatcacaaatttattttaaatgatatgcgctttgtgttattctcatatttaaatttcagTGGTCTTGCTGCCGGATTCCTTTTGCTAAAGTTGCAAAAGCAAAGAGAGAAAATGTACCACgtgagtgaaattagacacggttgaggtagatcacaagaccaacagggcttgacttacagGTGTTAGATCAAGGGTTGGGTAGGAATAGAAATGAAGAACATGACAGTCATCATTTCAAGCTGAATAAGTTATTTATATCAATCAGTTTTTCGTTTGCAGCTGCATTATCCTTTGTACAAATTGgtaacaaaaaagtatgactaattcgaaaccattcgaaaaatttccagCTACAAAAATGAGATATCTTGATCTCCCTGACACTCCTGATCTGATTAACAGAAAAATCTACTTTGTGATTCTTGAAGGATACGTGgtacggttgatattctttatgcttgccaaaaattttcaaaattaagATTTTCATCGCTTCCAGTTGCATTTCATTAGTCGCAATATTATTTTTTCAACTATGTAGAATACTCCAATTTTCGATAGGACAACTAATAGAAGAAATGGTGCCTAAAGAAAAAGAAGAATGCCCACTACCAGAACAATCTTTACAACAAATTCACGATGTTCAAATTCATTACCAAGAAATTTCCAATGCTAAGCTGTATATAGAACAGAATTTCTCATTTTCACTGTTTTATACCTACGGC**TCTTCA**ATTCCG**TTA**ACCTGTCTGTTGGGCTACATTGCTTTTCGAAATGGAATTCAGgtagtgaaaatgagcagcaataccattgtttcatatcatcatttagGCTGATATGGCGGAAACTTTTTCTGTGGCAATATGGCTCACTAATACTATGTTAGCATTGATGCTTTTCTCGATTCCAGCTTTTATGATTGCTGAAGAGgttagtacttataagtatgtatatgtttgtttaaatacatagatatagttagagcagaactgagaaaattgagttagtaaagtttttgtatttcttcaacttttgatgttagctgaacaaaaattaaaaaacaacataaattaaatataatgcaagcatatttgtgaacaagaaatgtgtacgtggataaagtcgcggtggaattatcatttaaccgtgattgcatgcatataaatcccacaatacgtaaactcaatttatttatgacagtttcacaattacgatcagaaatttaaaataaataggttttcaaacactattcttcagcgacaatatttctgctaattgtcgctgatattttatgttcgcaaacgcagtaattcaaaattgatatccaaacattaagttgtacaaataaaaattaataattacagGGCGACAAACTTCTAACCGCTTCTTTCAAAATGTATCACGAAACTCTATGTGAGGAACGTGATCTTCTAGTTTTGgtgtgttcgttatctcaaaaaaaaatcgatcaattctactttcagTCTCAAATGAGTTTTCTTTCGTTTCAAATGCATGCTACAAAGCTTACTCTCACCGCCGGGAATTTCTTCATGATGAATCGCAAAATTATGATCAGTgtacgtttaaaggcgtgcagctggtttttagttgatatttaaaatatttcagTTGTTCTCTGCAATTTTCACATATTTTCTTATTTTGGTACAGTTTGATGCTGAAAAGGAGCGAGCCGGAGAATGCAATAATCAATCTCGTGTGCTCATTGTCCAACCACCTGTGTAAataaaattgtcttaacattttccccatattgagccaacttcaattagtttccaatatgttcatttgtaactacgcaatgtatttgacaaattgtttattcatgtctcctaaatacaagaaagaaagtcattt

Missense mutations are labeled in **red**. Synonymous mutations are labeled in **blue**.

A Exon

A Exon

a Intron

a UTR
